# Altered PI3K-PTEN balance promotes preferential killing of human IgE^+^ plasma cells by BCR crosslinking

**DOI:** 10.64898/2026.05.28.728415

**Authors:** Faruk Ramadani, Helena Tolarova, Sin Wah Tooki Chu, Clare Thomas, Line Ohm-Laursen, Pavel Tolar

## Abstract

Immunoglobulin E (IgE) drives allergic disease, yet what restrains the persistence of IgE production remains poorly understood. Mouse studies suggest that BCR-induced apoptosis limits the survival of IgE-producing plasma cells (PCs). Whether this mechanism applies to human IgE⁺ PCs is unclear. Using a human IgE class-switching system, we show that BCR crosslinking preferentially kills IgE⁺ PCs compared to IgG1^+^ PCs. However, this selective sensitivity is not explained by surface BCR levels or proximal BCR signaling as suggested in mice. Instead, elevated PTEN expression in IgE⁺ PCs constrains PI3K/Akt pro-survival signaling and lowers the apoptotic threshold by upregulating BIM, while JNK signaling sustains PTEN expression and amplifies their apoptotic sensitivity. CRISPR/Cas9 targeting of PTEN or BIM, or JNK inhibition protects IgE⁺ PCs from BCR-mediated killing. Therapeutic anti-IgE antibodies, including omalizumab and extracellular membrane-proximal domain (EMPD)-targeting antibodies, exploit this sensitivity to selectively eliminate IgE⁺ PCs and suppress IgE production, providing a mechanistic rationale for depleting IgE⁺ PCs in allergic disease.

**Summary:** Ramadani et al. identify a JNK/PTEN/BIM signaling axis that intrinsically limits human IgE⁺ plasma cell survival and drives their preferential sensitivity to BCR-induced apoptosis. This mechanism is distinct from that established in mice and has direct implications for anti-IgE therapeutic strategies.

## Introduction

Immunoglobulin E (IgE) plays an important role in protective immunity against parasites but is also the principal driver of allergic disease (Rahman and Wesemann, 2024; Dullaers et al., 2012). As a result, IgE production is tightly regulated compared to other antibody isotypes, reflecting the need to balance protective functions against the risk of allergic responses. A hallmark of this regulation is the absence of canonical IgE^+^ memory B cells and the limited persistence of IgE⁺ plasma cells (PCs), which contrasts with the long-lived PC populations that sustain IgM, IgG and IgA responses (Yang et al., 2012; Talay et al., 2012a; Haniuda et al., 2016; Talay et al., 2012b; He et al., 2013). Indeed, IgE⁺ PCs are characterised by high secretory activity and elevated endoplasmic reticulum (ER)stress (Vecchione et al., 2024), features that could be contributing to their reduced lifespan. However, recent studies have also identified populations of long-lived IgE^+^ PCs in various tissues (Asrat et al., 2020; Vecchione et al., 2024; Ding et al., 2025; Miranda-Waldetario et al., 2025) demonstrating that IgE^+^ PCs can persist under certain conditions. These observations raise the possibility that IgE⁺ PCs exist near a critical threshold between survival and apoptosis, but the molecular mechanisms that regulate this balance remain unclear.

Several lines of evidence indicate that the IgE B cell receptor (BCR) is a key regulator of IgE^+^ cell fates. Unlike the BCR of other isotypes, the IgE BCR can constitutively signal independently of antigen binding (Newman and Tolar, 2021; Yang et al., 2016; Haniuda et al., 2016). This contributes to the precocious differentiation of IgE^+^ GC B cells into PCs via the PI3K-IRF4 pathway (He et al., 2013; Erazo et al., 2007; Yang et al., 2012; Newman and Tolar, 2021; Haniuda et al., 2016; Yang et al., 2016). In parallel, this constitutive antigen-independent signaling activates PLCγ2 and sustains intracellular Ca²⁺ flux, which has been linked to the increased apoptotic susceptibility of IgE⁺ PCs through induction of pro-apoptotic of Bcl2-family members, including BIM(Newman and Tolar, 2021). Furthermore, genetic defects in BCR signaling have been linked to enhanced and sustained IgE production (SoRelle et al., 2021; Haniuda et al., 2016). Together, these observations suggest that the IgE BCR couples differentiation and cell death programmes, thereby limiting IgE production.

Consistent with this, studies in murine systems have shown that compared to other PCs, IgE⁺ PCs are more sensitive to antigen crosslinking of their BCR in inducing apoptosis, and this results in their preferential killing both in vitro and in vivo (Wade-Vallance et al., 2023). This susceptibility to apoptosis by BCR crosslinking has been attributed to increased surface expression of the IgE BCR on IgE^+^ PCs and enhanced activation of proximal BCR signaling, such as Syk and PLCγ2, leading to elevated Ca²⁺ flux (Wade-Vallance et al., 2023). However, this model does not fully explain isotype-specific differences in apoptotic susceptibility. For example, mouse IgM^+^ PCs have similar levels of surface BCR and proximal BCR signaling but are resistant to the apoptosis induced by BCR crosslinking (Wade-Vallance et al., 2023). This indicates that additional factors beyond proximal signaling strength may influence the outcome of BCR crosslinking across PCs of different Ig isotypes. Furthermore, whether this mechanism applies to human IgE⁺ PCs has yet to be determined. This is particularly important because human IgE-switched cells express two isoforms of membrane IgE (mIgE), a short and long form, which differ in their surface expression and signaling capacity, and are differentially regulated during PC differentiation (Ramadani et al., 2017; Batista et al., 1996; Vanshylla et al., 2018).

These mechanistic questions are also directly relevant to the therapeutic targeting of IgE responses. Omalizumab, which is widely used to treat allergic disease, primarily neutralises circulating IgE and blocks its interaction with Fc ε receptors (Eggel et al., 2024; Guntern and Eggel, 2020; Holgate, 2014). However, as omalizumab also binds mIgE (Ramadani et al., 2017; Chen et al., 2010), it has the potential to directly crosslink the IgE BCR and modulate the survival of IgE^+^ PCs. Antibodies targeting the extracellular membrane-proximal domain (EMPD) of human mIgE provide a complementary strategy to selectively crosslink the IgE BCR independently of soluble IgE (Chen et al., 2010; Liour et al., 2016; Feichtner et al., 2008; Brightbill et al., 2010). Whether such interventions suppress IgE production partly by inducing the apoptosis of IgE⁺ PCs remains to be determined.

In this study, using a human tonsil-derived class-switching system, we show that BCR crosslinking preferentially induces apoptosis in human IgE⁺ PCs compared with IgG1⁺ PCs. Mechanistically, we identify an imbalance between survival and stress signaling pathways as the key determinant of this sensitivity. Specifically, elevated PTEN expression in IgE⁺ PCs constrains PI3K/Akt signaling, thereby lowering the apoptotic threshold by leading to higher induction of the pro-apoptotic protein BIM. In parallel, JNK signaling acts as a critical amplifier of this response, reinforcing PTEN expression and promoting BIM-dependent apoptosis. Consistent with this model, we further show that therapeutic anti-IgE antibodies, including omalizumab and EMPD-targeting antibodies, suppress IgE production by selectively eliminating IgE⁺ PCs through BCR crosslinking. These findings define the altered balance between PI3K and a coordinated JNK/PTEN//BIM axes that predisposes human IgE⁺ PCs to apoptosis and provide mechanistic insights for understanding how IgE production is intrinsically limited and may be therapeutically targeted.

## Results

### Omalizumab and anti-CεmX antibodies reduce IgE production by killing IgE-producing cells

To assess the ability of anti-IgE antibodies to eliminate IgE⁺ cells, we used our previously characterised human tonsil culture system for IgE class switching induced by IL4 and anti-CD40(Ramadani et al., 2019, 2015, 2017). After 12 days of culture, class-switched cells were harvested and cultured for a further 48 h either in medium alone or in the presence of anti-IgE antibodies. These included omalizumab, a therapeutic antibody that binds the CH3 region of membrane IgE (mIgE) (Holgate, 2014; Eggel et al., 2024), and anti-IgE CεmX 4B12, which targets the extracellular membrane-proximal domain (EMPD) of the long form of mIgE (Chen et al., 2010). As a control, we used anti-CεmX a20, which cannot bind its epitope within the EMPD because this region is masked by Igα and Igβ (Chen et al., 2010).

As expected, anti-IgE CεmX a20 had no effect on the number of IgE^+^ cells in cultures (Fig. 1A and 1B). In contrast, stimulation with anti-IgE CεmX 4B12 and omalizumab significantly reduced the numbers of IgE^+^ cells compared with control cultures (Fig. 1A and 1B). However, anti-CεmX 4B12 reduced IgE⁺ cells more efficiently (down to 52%±12% of control) than omalizumab (down to 81.2%±9.5% of control) (Fig. 1A and 1B). The proportions of IgG1^+^ cells remained unchanged in both anti-CεmX 4B12 and omalizumab stimulated cultures, confirming that the effect of these antibodies is specific to IgE^+^ cells (Fig. 1A and 1B).

**Figure 1.**
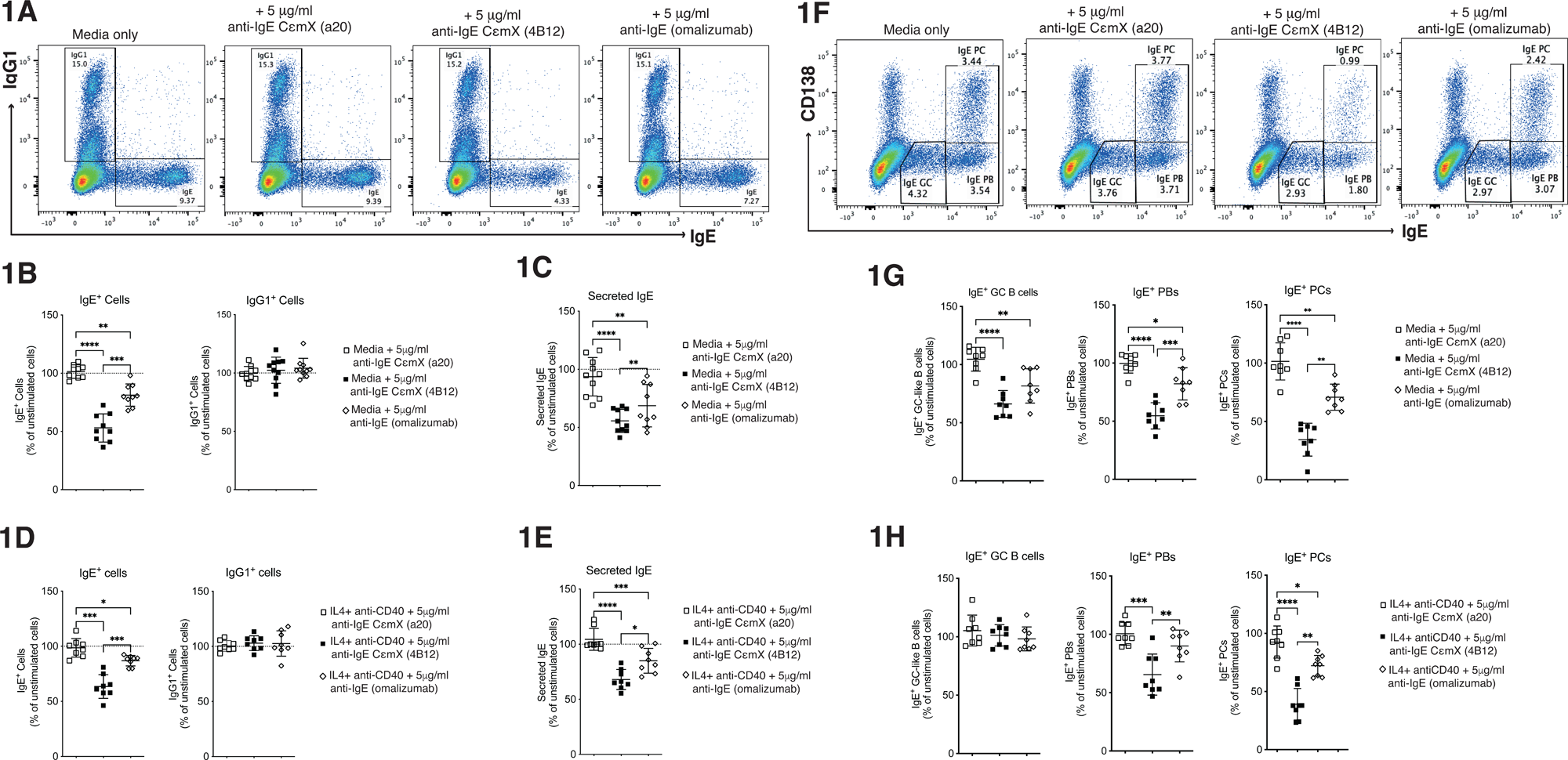
Anti-IgE antibodies selectively reduce IgE⁺ cells and IgE secretion. (A) Representative flow cytometry plots showing IgE⁺ and IgG1⁺ cells cultured in media alone or in the presence of 5 μg/mL anti-IgE antibodies (CεmX clones a20 and 4B12, or omalizumab). (B) IgE⁺ and IgG1⁺ cells following stimulation with 5 μg/mL anti-IgE antibodies, shown as a percentage of unstimulated media alone culture. (C) Secreted IgE levels in culture supernatants following stimulation with anti-IgE antibodies, shown as a percentage of unstimulated media alone culture. (D) IgE⁺ and IgG1⁺ cells cultured with IL-4 and anti-CD40 in the presence of 5 μg/mL anti-IgE antibodies. Data shown as a percentage of IL-4 and anti-CD40 culture. (E) Secreted IgE levels under IL-4 and anti-CD40 stimulation in the presence of 5 μg/mL anti-IgE antibodies. Data shown as a percentage of IL-4 and anti-CD40 culture. (F) Representative flow cytometry plots showing differentiation of IgE⁺ cells into GC-like B cells (IgE^low^CD138^-^), plasmablasts (PBs; IgE^hi^CD138^-^), and plasma cells (PCs; IgE^hi^CD138⁺), in the presence of media alone or 5 μg/mL of anti-IgE antibodies (CεmX clones a20 and 4B12, or omalizumab). (G) IgE⁺ GC-like B cells, PBs, and PCs following stimulation with 5 μg/mL anti-IgE antibodies, shown as a percentage of unstimulated media alone culture. (H) IgE⁺ GC-like B cells, PBs, and PCs cultured with IL-4 and anti-CD40 and in the presence of 5 μg/mL anti-IgE antibodies. Data is shown as a percentage of unstimulated IL-4 and anti-CD40 culture. Data are pooled from independent experiments (symbols represent individual donors). Data indicate mean ± SD. Statistical significance was determined using one-way ANOVA with Tukey’s multiple comparison test. *P < 0.05, **P < 0.01, ***P < 0.001, ****P < 0.0001.

Because IgE⁺ cells arise in environments that stimulate IL-4 receptor and CD40 signaling, which can protect B cells from the BCR-induced apoptosis (Tsubata et al., 1993; Parry et al., 1994; Berard et al., 1999), we next examined whether continued presence of these signals can rescue IgE^+^ cells from the effects of mIgE crosslinking. Figure 1D shows that even in the presence of IL-4 and anti-CD40 the proportions of IgE^+^ cells, although not as pronounced, were still significantly reduced by either anti-IgE CεmX 4B12 and omalizumab. Thus, crosslinking of the mIgE reduces the numbers of IgE^+^ cells regardless of co-stimulation through IL-4 receptor and CD40.

Consistently, we found that adding either anti-CεmX 4B12 or omalizumab to the cultures significantly lowered the amounts of secreted IgE in supernatants compared to controls (Fig. 1C and E). Together, these data indicate that crosslinking of mIgE with either omalizumab or anti-CεmX 4B12 suppresses IgE production by selectively reducing the number of IgE^+^ cells.

We previously reported that differentiation of human IgE⁺ B cells into IgE⁺ PCs is accompanied by increased sensitivity to apoptosis (Ramadani et al., 2017). Similar observations were also made in studies of mouse IgE^+^ cells (Haniuda et al., 2016; Newman and Tolar, 2021). When examining the effect of mIgE crosslinking on different stages of IgE⁺ cell differentiation, we found that crosslinking mIgE with anti-CεmX 4B12 or omalizumab significantly reduced IgE⁺ cells across all stages of differentiation compared to cultures maintained in medium alone (Fig. 1F and G). However, IgE⁺ GC-like B cells were the least affected, while IgE⁺ PCs were the most affected, indicating that IgE^+^ PCs are particularly sensitive to mIgE crosslinking (Fig. 1G). This reduction in cell numbers correlated with increased cell death, indicating that mIgE crosslinking induces apoptosis of IgE⁺ cells, particularly at the PCs stage (Fig. S1).

In addition, IgE⁺ PB and PCs remained sensitive to mIgE crosslinking in the presence of IL-4 and anti-CD40 under these conditions, whereas IgE⁺ GC B cells were rescued by the co-stimulations (Fig. 1H and Fig. S1D), indicating that IL-4 and CD40 signaling selectively protect the less differentiated IgE⁺ cells, whereas more differentiated IgE⁺ cells are susceptible to apoptosis induced by anti-CεmX 4B12 and omalizumab.

### BCR crosslinking kills human IgE^+^ PC more efficiently than IgG1^+^ PCs

It was recently reported that mIg crosslinking kills mouse IgE⁺ PCs more efficiently than IgG1⁺ PCs (Wade-Vallance et al., 2023). To determine whether this is also the case in the human system, we used anti-λ and anti-κ F(ab’)₂ antibodies to crosslink the BCR of cultured IgE⁺ and IgG1⁺ cells.

As shown in Figure 2A and 2B, IgE^+^ and IgG1^+^ GC-like B cells were unaffected by anti-λ and anti-κ F(ab)^2^ stimulations compared with control cultures. In contrast, the proportion of both IgE^+^ and IgG1^+^ plasmablasts (PBs) were reduced at similar rates following stimulation with anti-λ and anti-κ F(ab)^2^ antibodies. However, IgE^+^ PCs were significantly more sensitive to BCR crosslinking than IgG1^+^ PCs, suggesting that at the PC stage of differentiation IgE^+^ cells are preferentially killed by BCR crosslinking compared to IgG1^+^ cells (Figure 2A and 2B).

**Figure 2.**
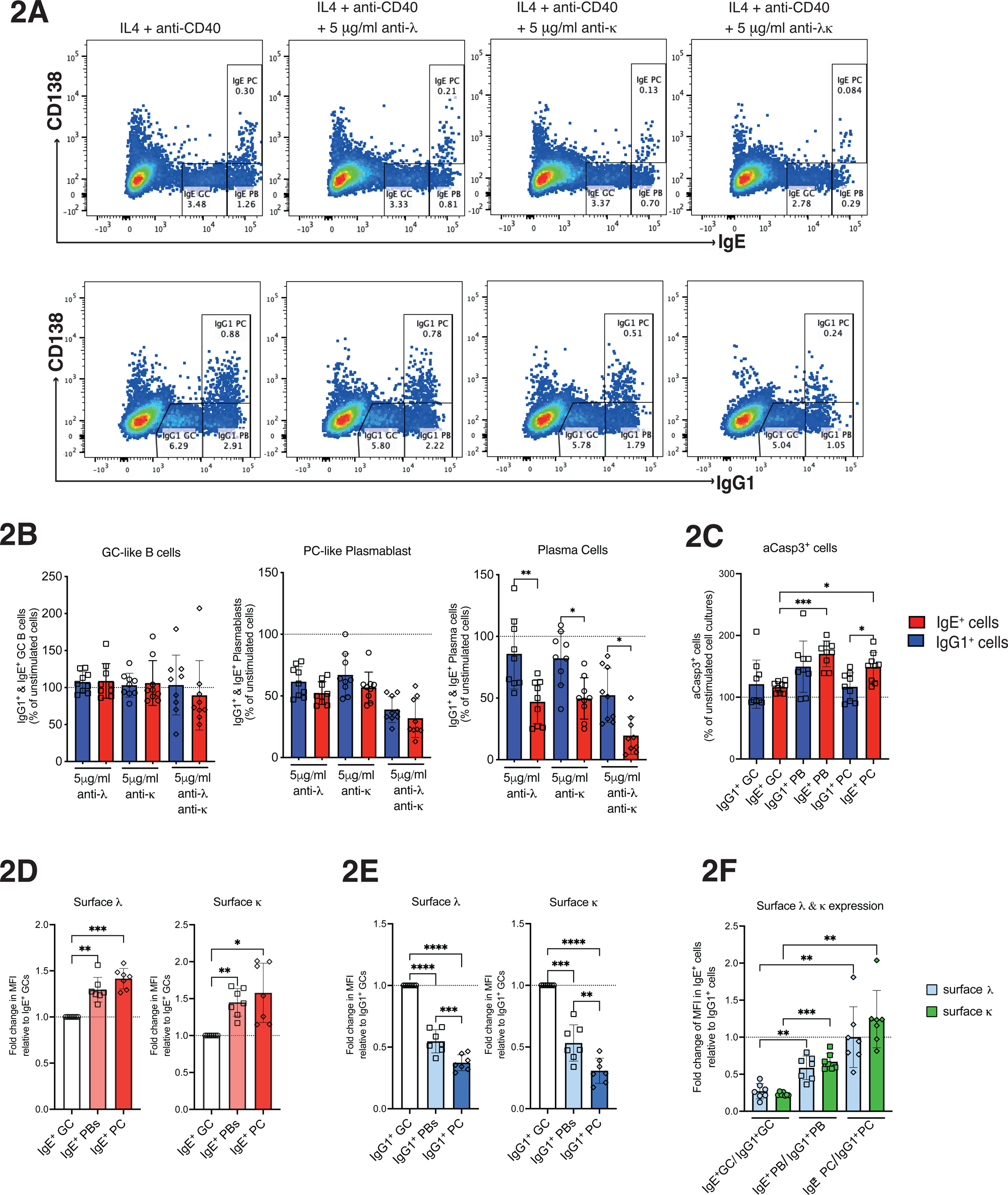
BCR crosslinking selectively induces apoptosis in IgE⁺ PCs. (A) Representative flow cytometry plots of IgE⁺ (top) and IgG1⁺ (bottom) cells following culture with IL-4 and anti-CD40, with or without BCR crosslinking using (5 μg/mL) F(ab’)_2_ anti-λ, (5 μg/mL) F(ab’)_2_ anti-κ or (5 μg/mL) combined F(ab’)_2_ anti-λ/κ antibodies. GC-like B cells, PBs, and PCs are defined by their CD138 and Ig expression. (B) IgE^+^ and IgG1⁺ GC-like B cells, PBs, and PCs following BCR crosslinking with F(ab’)_2_ anti-λ, F(ab’)_2_ anti-κ or combined F(ab’)_2_ anti-λ/κ antibodies, expressed as a percentage of unstimulated 48 h IL-4 and anti-CD40 culture. (C) Quantification of active caspase-3⁺ within IgE⁺ and IgG1⁺ populations across GC-like B cells, PBs and PCs following 1 h of BCR crosslinking. The data are shown as a percentage of unstimulated IL-4 and anti-CD40 culture. (D) Quantification of surface λ and κ mean fluorescence intensity (MFI) across differentiation stages of IgE⁺ cells made relative to the MFI of IgE^+^ GC-like B cells. (E) Quantification of surface λ and κ mean fluorescence intensity (MFI) across differentiation stages of IgG1⁺ cells made relative to the MFI of IgG1^+^ GC-like B cells. (F) Comparison of total surface BCR (λ + κ) expression between IgE^+^ and IgG1^+^ cells. The data show the fold change in MFI surface λ and κ on IgE^+^ cell populations made relative to their respective IgG1^+^ cell counterparts. Data are pooled from independent experiments (symbols represent individual donors). Data indicate mean ± SD. Statistical significance was determined using one-way ANOVA with Tukey’s multiple comparison test. *P < 0.05, **P < 0.01, ***P < 0.001, ****P < 0.0001.

To confirm the increased apoptosis of IgE^+^ PCs, cultured cells were stimulated for 1 hour with anti-λ and anti-κ F(ab)^2^ antibodies and the levels of active Caspase 3 were measured (Figure 2C). Following stimulation, active caspase-3 levels were similar between IgE^+^ and IgG1^+^ GC-like B cells. However, active caspase-3 levels increased amongst the PBs and PCs with a significant difference between IgE^+^ and IgG1^+^ cells at the PC stage. Together, these data indicate that BCR crosslinking by anti-λ and anti-κ F(ab)^2^ antibodies preferentially kills human IgE^+^ PCs compared to IgG1^+^ PCs.

To determine whether differences in BCR expression contribute to the selective sensitivity of IgE⁺ PCs to BCR crosslinking, we quantified surface BCR expression across IgE^+^ and IgG1^+^ cells in all differentiation stages by staining for surface λ and κ light chains. Consistent with our previous studies(Ramadani et al., 2017), surface λ and κ expression increased as IgE⁺ cells differentiated into PCs, whereas both were downregulated during IgG1^+^ PC differentiation (Figure 2D and 2E). Despite this, comparison between IgE^+^ and IgG1^+^ cells revealed that while BCR expression at the GC-like stage was approximately fourfold lower in IgE^+^ relative to IgG1^+^ cells at the PC stage IgE⁺ cells expressed surface BCR at levels similar to IgG1⁺ cells (Figure 2F). This finding contrasts with previous observations in mice, where IgE^+^ PCs express higher surface BCR levels relative to other PC subsets (Wade-Vallance et al., 2023), and indicates that differences in surface BCR expression do not account for the preferential killing of human IgE^+^ PCs by BCR crosslinking.

### Syk and PLCγ2 signaling contribute to BCR killing of PCs but do not account for the preferential killing of IgE⁺ PCs

Previous work in mice demonstrated that BCR crosslinking induces stronger proximal BCR signaling in IgE⁺ PCs than in IgG1⁺ PCs, leading to their apoptosis, and that inhibition of Syk or targeting of PLCγ2 rescues IgE⁺ PCs from this BCR-mediated killing (Wade-Vallance et al., 2023). To determine whether a similar mechanism operates in human IgE^+^ PCs, we first examined the phosphorylation of Syk along the differentiation pathway of human IgE^+^ and IgG1^+^ B cells.

Basal Syk phosphorylation was comparable between IgE^+^ and IgG1^+^ cells (Figure 3A). However, following 1 and 10 min of BCR crosslinking, IgG1^+^ GC-like B cells showed higher Syk phosphorylation than IgE^+^ GC-like B cells (Figure 3B and Figure S2A). In contrast, at the PB and PC stages of differentiation, Syk phosphorylation was similar between IgE^+^ and IgG1^+^ cells (Figure 3B and Figure S2B). To assess the contribution of Syk signaling to BCR-mediated killing of IgE^+^ and IgG1^+^ PCs, day 12 class-switched cells were cultured for a further 48 h with three different concentrations of Syk inhibitor PRT062607 Hydrochloride (250 nM, 1 μM, or 4 μM), in the presence or absence of BCR crosslinking.

**Figure 3.**
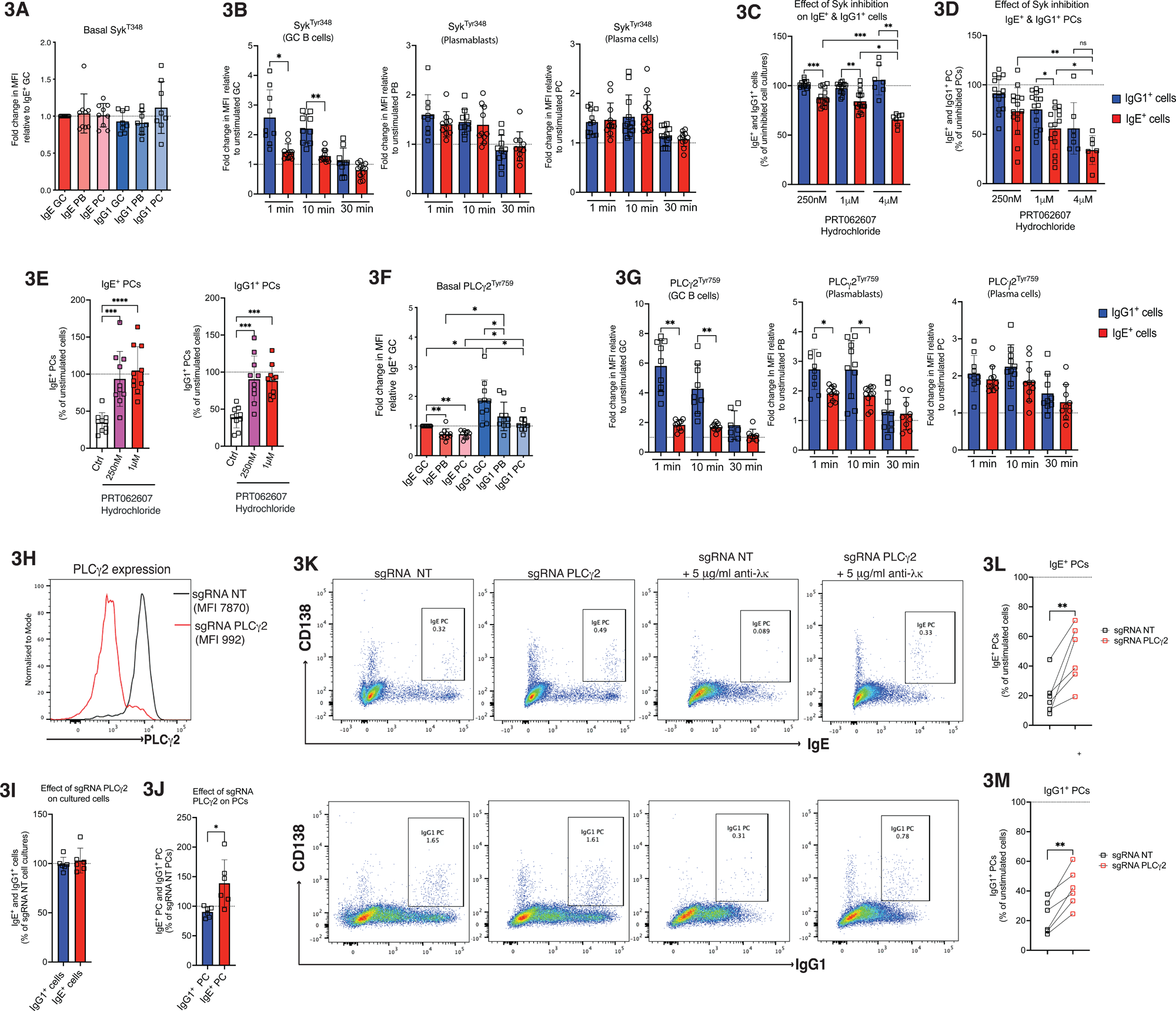
Syk inhibition and CRISPR/Cas9 PLCγ2 targeting protect IgE^+^ and IgG1^+^ PCs from BCR mediated killing. (A) Basal phosphorylation of Syk (Tyr348) in IgE⁺ and IgG1⁺ cells across GC-like B cells, plasmablasts (PBs), and plasma cells (PCs). Fold change in MFI is shown relative to IgE⁺ GC-like B cells. (B) Syk phosphorylation following BCR crosslinking, at different timepoints, with F(ab’)_2_ anti-λ/κ (10 μg/mL) in GC-like B cells, PBs, and PCs. IgG1⁺ cells (blue) IgE⁺ cells (red). The data show the fold change in MFI relative to unstimulated GC-like B cells, PBs, and PCs. (C) Effect of Syk inhibition (PRT062607 hydrochloride) on IgE⁺ and IgG1⁺ cells in the 48 h cultures. The data are shown as a percentage of uninhibited cell culture. (D) Effect of Syk inhibition on IgE^+^ and IgG1^+^ PCs gated within the total IgE^+^ and IgG1^+^ cells. The data are shown as a percentage of uninhibited cell culture. (E) IgE⁺ (left) and IgG1⁺ (right) PCs following 48 h of stimulation with 5 μg/ml of F(ab′)₂ anti-λ/anti-κ in the presence of two different concentrations of Syk inhibitor. Ctrl = IL4 + anti-CD40 with DMSO. Data is shown as a percentage of the unstimulated cell culture. (F) Basal phosphorylation of PLCγ2 (Tyr759) in IgE⁺ and IgG1⁺ cells across GC-like B cells, plasmablasts (PBs), and plasma cells (PCs). Fold change in MFI is shown relative to IgE⁺ GC-like B cells. (G) PLCγ2 phosphorylation following BCR crosslinking with F(ab’)_2_ anti-λ/κ (10 μg/mL) in GC-like B cells, PBs, and PCs. IgG1⁺ cells (blue) IgE⁺ cells (red). The data show the fold change in MFI relative to unstimulated GC-like B cells, PBs, and PCs. (H) Representative histogram confirming the successful PLCγ2 targeting by CRISPR/Cas9 (sgRNA PLCγ2) compared with non-targeting control (sgRNA NT). (I) Effect of PLCγ2 targeting on overall IgE⁺ and IgG1⁺ cell proportions in class switching cultures. The data are shown as a percentage of sgRNA NT cell culture. (J) Effect of PLCγ2 targeting on IgE^+^ and IgG1^+^ PCs gated within the total IgE^+^ and IgG1^+^ cells. The data are shown as a percentage of sgRNA NT cell culture. (K) Representative flow cytometry plots showing IgE⁺ PCs (top row) and IgG1^+^ PCs (bottom row) in sgRNA NT and sgRNA PLCγ2 48 h cultures, with or without 5 μg/ml of F(ab’)_2_ anti-λ/κ antibodies. (L) IgE^+^ PCs and (M) IgG1⁺ PCs in sgRNA NT and sgRNA PLCγ2 cell cultures following BCR crosslinking. Data is shown as a percentage of unstimulated culture. Each symbol represents individual donors. The data indicate mean ± SD. Statistical significance was determined using one-way ANOVA with Tukey’s multiple comparison test (B, C, D, E, F, G) or paired two-tailed t-test with Welch’s correction (J, L, M). *P < 0.05, **P < 0.01, ***P < 0.001.

In the absence of BCR-crosslinking, Syk inhibition reduced the proportion of IgE⁺ cells across all concentrations tested, whereas the proportions of IgG1⁺ cells remained unchanged (Figure 3C), suggesting that Syk inhibition selectively affects IgE^+^ cells in these cultures. Analysis of the PCs within the gated IgE^+^ and IgG1^+^ cells showed that the high dose Syk inhibition (4 μM) significantly reduced both IgE^+^ and IgG1^+^ PCs, whereas the lower concentrations (250 nM and 1 μM) led to a slight reduction, which was more pronounced in IgE^+^ PCs (Figure 3D). Considering this, subsequent analyses focused on the effects of 250 nM and 1 μM PRT062607 Hydrochloride on BCR-mediated killing. Under these conditions, Syk inhibition rescued both IgE^+^ and IgG1^+^ PCs from BCR-mediated killing (Figure 3E, Figure S2C and S2D). A similar protective effect of Syk inhibition was also observed in IgM⁺ PCs (Figures S2E–H). IgM⁺ cells maintain their BCR expression upon PC differentiation, resulting in higher surface BCR levels than those observed on IgG1⁺ and IgE⁺ PCs (Figure S2I and S2J). However, the low frequency of IgM⁺ PCs detectable in our cultures prevented the measurement of BCR signaling in these cells. Nevertheless, the shared dependence on Syk activity of IgE⁺, IgG1⁺ and IgM⁺ PCs indicates that BCR signaling through Syk is a common requirement for BCR-mediated PC killing, irrespective of Ig isotype or surface BCR expression level.

We next examined the PLCγ2 activation and found that IgG1^+^ GC-like B cells displayed the highest basal PLCγ2 phosphorylation, which significantly reduced upon PC differentiation (Figure 3F and Figure S2B). A similar trend was observed for IgE^+^ cells, which overall had significantly lower basal PLCγ2 phosphorylation than IgG1^+^ cells (Figure 3F and Figure S2B). Following 1 and 10 min of BCR crosslinking, the PLCγ2 phosphorylation remained significantly higher in IgG1^+^ GC-like B cells and PBs compared with IgE^+^ cells (Figure 3G and Figure S2B). However, at the PC stage of differentiation, the PLCγ2 phosphorylation was only slightly but not significantly higher in IgG1^+^ compared to IgE^+^ and cells (Figure 3G). To assess the role of PLCγ2 in the BCR-mediated killing of IgE^+^ PCs, we successfully targeted the gene encoding PLCγ2 using a CRISPR/Cas9 electroporation approach (Figure 3H). The PLCγ2 targeting did not alter the overall proportions of IgE^+^ and IgG1^+^ cells (Figure 3I) but increased the proportions of IgE^+^ PCs (Figure 3J), which is consistent with the role of this pathway in promoting apoptosis and inhibiting differentiation of IgE^+^ PCs(Newman and Tolar, 2021). Importantly, as in the case of Syk inhibition, PLCγ2 targeting rescued the BCR-mediated killing of both IgE^+^ and IgG1^+^ PCs (Figure 3K-M).

Because BCR activation of PLCγ2 is required for Ca^2+^ mobilisation via the inositol trisphosphate receptor (IP3R)(Baba et al., 2014; Saleem et al., 2014), we next assessed the contribution of Ca^2+^ flux in BCR-mediated killing of IgE^+^ PCs using the IP3R antagonist 2-aminoethoxydiphenyl borate (2-APB)(Baba et al., 2014; Saleem et al., 2014). Day 12 class-switched cells were cultured for a further 48 h with two different concentrations of 2-APB (25 μM and 50 μM), in the presence or absence of BCR crosslinking. Inhibition of IP3R Ca^2+^ release significantly reduced the BCR-mediated killing of IgE^+^ PCs (Figure S3 A and C), and it also protected IgG1^+^PC but to a lesser extent than IgE^+^ PCs (Figure S3 B and D), indicating that Ca2^+^ flux is a key downstream effector of the BCR-mediated killing of human PCs, and it contributes to the greater sensitivity of IgE^+^ PCs.

Together, these data indicate that Syk, PLCγ2 and downstream Ca2^+^ signaling contribute to the BCR-mediated killing of human PCs. However, in contrast to the mouse system, Syk and PLCγ2 signaling and BCR surface levels do not differ between IgE^+^ and IgG1^+^ PCs suggesting that the increased susceptibility of human IgE^+^ PCs to BCR-mediated killing relies on different mechanisms.

### Elevated PTEN expression predisposes IgE^+^ PCs to BCR-mediated apoptosis

Given the central role of PI3K signaling in BCR-dependent B cell survival (Werner et al., 2010; Srinivasan et al., 2009), we assessed the activity of this pathway in IgE^+^ and IgG1^+^ cells. While the basal Akt phosphorylation was similar between IgE^+^ and IgG1^+^ cells (Figure 4A and Figure S4A), BCR crosslinking for 1 and 10 mins induced significantly higher Akt phosphorylation in IgG1^+^ GC-like B cells and PBs compared to IgE^+^ cells (Figure 4B and Figure S4A). Notably, this difference persisted in the PCs, with IgG1^+^ PCs showing higher Akt phosphorylation than IgE^+^ PCs after stimulation (Figure 4B and Figure S4A), indicating that the activation of PI3K signaling downstream of the BCR in IgE^+^ PC is impaired. To understand the basis of this reduced Akt phosphorylation, we measured the levels of PTEN, a phosphatase that negatively regulates PI3K signaling (Okkenhaug and Vanhaesebroeck, 2003). These experiments revealed that IgE^+^ PCs express significantly higher levels of PTEN than IgG1^+^ and IgM^+^ PCs (Figure 4C and 4D), suggesting that the elevated PTEN may limit the PI3K-dependent survival signals in IgE^+^ PCs. To test this, we targeted PTEN using CRISPR/Cas9 electroporation at the onset of the culture (Figure 4E). PTEN deficiency increased the overall cell viability of our cultures (Figure 4F) but drastically reduced the proportion of total IgE^+^ cells (Figure 4G and Figure S4B), which is consistent with previous reports that PTEN deficiency inhibits class switching to IgE (Chen et al., 2015). To overcome this effect, we targeted PTEN on day 7 of the culture, after class switching had been initiated. Under these conditions, PTEN targeting similarly enhanced cell viability of the culture (Figure 4F), while its impact on IgE class switching was substantially lower compared to targeting at the onset of the culture (Figure 4G and Figure S4C). Notably, cultures lacking PTEN contained a 5- to 7-fold increase in IgG1^+^ PCs compared with control cultures electroporated with non-targeting sgRNA (Figure 4H, S4B and S4C), indicating that PTEN constrains IgG1^+^ PC accumulation. Furthermore, day 7 PTEN targeting also increased the proportions of IgM^+^ PCs (Figure S4D) and to some degree IgE^+^ PCs (Figure S4D). These results are consistent with previous reports showing that the PI3K pathway promotes PC differentiation (Newman and Tolar, 2021; Cutrina-Pons et al., 2023).

**Figure 4.**
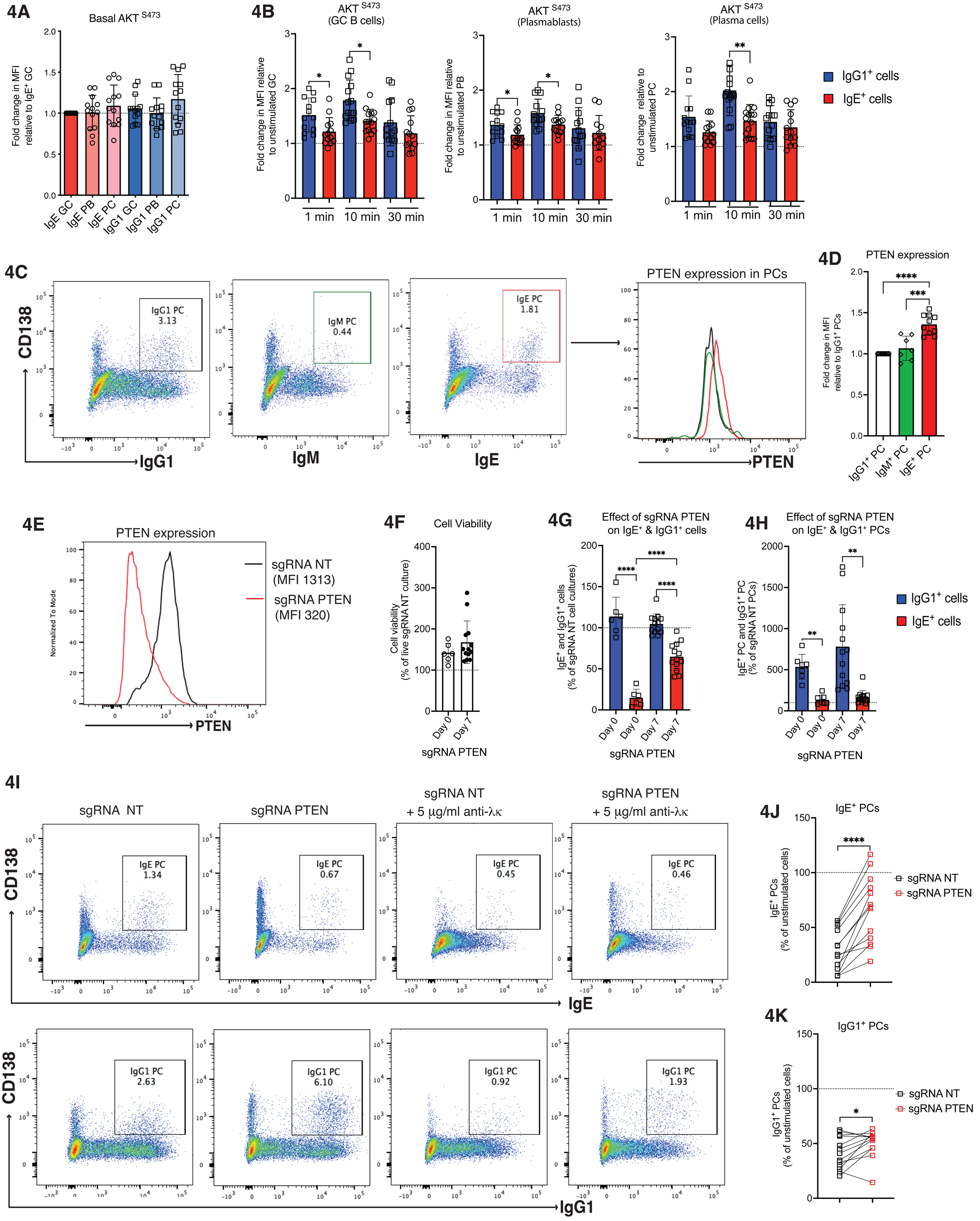
Elevated PTEN expression constrains PI3K–Akt signaling and promotes preferential BCR killing of IgE⁺ plasma cells. (A) Basal phosphorylation of Akt (Ser473) in IgE⁺ and IgG1⁺ cells across their PC differentiation pathway. Fold change in MFI is shown relative to IgE⁺ GC-like B cells. (B) Akt phosphorylation following BCR crosslinking with F(ab’)_2_ anti-λ/κ (10 μg/mL) in GC-like B cells, PBs, and PCs. IgG1⁺ cells (blue) IgE⁺ cells (red). The data show the fold change in MFI relative to unstimulated GC-like B cells, PBs, and PCs. (C) Representative flow cytometry plots showing identification of IgG1⁺, IgM⁺, and IgE⁺ PCs and corresponding PTEN expression histograms. (D) PTEN expression in IgG1⁺, IgM⁺ and IgE⁺ PCs. Data show the fold change in MFI relative to IgG1⁺ PCs. (E) Representative histogram confirming successful PTEN targeting by CRISPR/Cas9 (sgRNA PTEN) compared with non-targeting control (sgRNA NT). (F) Cell viability following PTEN targeting at day 0 and day 7 of culture, shown as a percentage of sgRNA NT cell cultures. (G) Effect of early (day 0) and delayed (day 7) PTEN targeting on IgE⁺ and IgG1⁺ cell proportions in class switching cultures. The data are shown as a percentage of sgRNA NT cell culture. (H) Effect of PTEN targeting on IgE^+^ and IgG1^+^ PCs gated within the total IgE^+^ and IgG1^+^ cells. The data are shown as a percentage of sgRNA NT cell culture. (I) Representative flow cytometry plots showing IgE⁺ PCs (top) and IgG1⁺ PCs (bottom) in sgRNA NT and sgRNA PTEN 48 h cultures, with or without 5 μg/ml of F(ab’)_2_ anti-λ/κ antibodies. (J) IgE⁺ PCs and IgG1⁺ PCs (K) in sgRNA NT and sgRNA PTEN cell cultures following BCR crosslinking. Data is shown as a percentage of unstimulated culture. Each symbol represents individual donors. The data indicate mean ± SD. Statistical significance was determined using one-way ANOVA with Tukey’s multiple comparison test (B, D, E, G, H) or paired two-tailed t-test with Welch’s correction (J, K). *P < 0.05, **P < 0.01, ***P < 0.001, ****P < 0.0001.

We next examined the role of PTEN in BCR-mediating killing and found that targeting PTEN led to a strong protection of IgE^+^ PCs from killing, whereas IgG1^+^ PCs were protected to a much lesser extent (Figure 4I-K). Furthermore, IgM^+^ PCs were not rescued by PTEN targeting (Figure SD and S4E). Together, these data indicate that the selectively elevated PTEN expression in IgE^+^ PCs limits their PI3K signaling, thereby weakening the survival signals and lowering the threshold for BCR-mediated killing of these cells.

### Attenuated PI3K signaling is associated with increased BIM induction and apoptosis in IgE^+^ PCs

We previously showed that inhibition of PI3K p110δ in these class switching cultures reduces the proportion of human IgE^+^ cells and blocks their differentiation into PCs (Cutrina-Pons et al., 2023). Consistent with this, inhibition of p110δ at day 10 for 48 h reduced IgE^+^ cells in cultures (Figure S5A and S5C), and impaired IgE^+^ and IgG1^+^ PC differentiation, as evidenced by the increased proportions of IgE^+^ and IgG1^+^ GC-like B cells and reduced proportions of IgE^+^ and IgG1^+^ PCs (Figure S5B and S5D). However, IgE^+^ PC were affected more strongly than IgG1^+^ PC, suggesting an additional mechanism beyond their impaired differentiation limits their accumulation.

Given that PI3K signaling promotes survival in part through suppression of pro-apoptotic factors(Duronio, 2008), we examined the expression of BIM protein, which has previously been linked to the increased apoptosis in IgE⁺ PCs (Ramadani et al., 2019; Newman and Tolar, 2021). Our data showed that inhibition of p110δ significantly upregulated BIM expression in IgE^+^ PCs but had minimal effect in IgG1^+^ PCs (Figure S5E and S5F), indicating that IgE^+^ PCs are more sensitive to the loss of PI3K-dependent survival signaling than IgG1^+^ PCs.

In class-switching cultures, BIM protein expression was comparable between IgE^+^ and IgG1^+^ PC (Figure 5A). However, BCR crosslinking induced a significantly greater increase in BIM expression in IgE^+^ PCs than in IgG1^+^ PCs (Figure 5A), suggesting that BCR crosslinking preferentially couples to pro-apoptotic pathways in IgE^+^ PCs. Given that PTEN limits PI3K signaling in IgE^+^ PCs, we investigated whether PTEN modulates BIM expression. PTEN targeting prevented the upregulation of BIM in IgE^+^ PCs following BCR crosslinking (Figure 5B and 5C), indicating that elevated PTEN expression in IgE^+^ PC promotes BIM expression by limiting the PI3K signaling. In contrast, PTEN targeting did not significantly change the levels of BIM expression in IgG1^+^ PCs following BCR crosslinking (Figure 5B and 5C).

**Figure 5.**
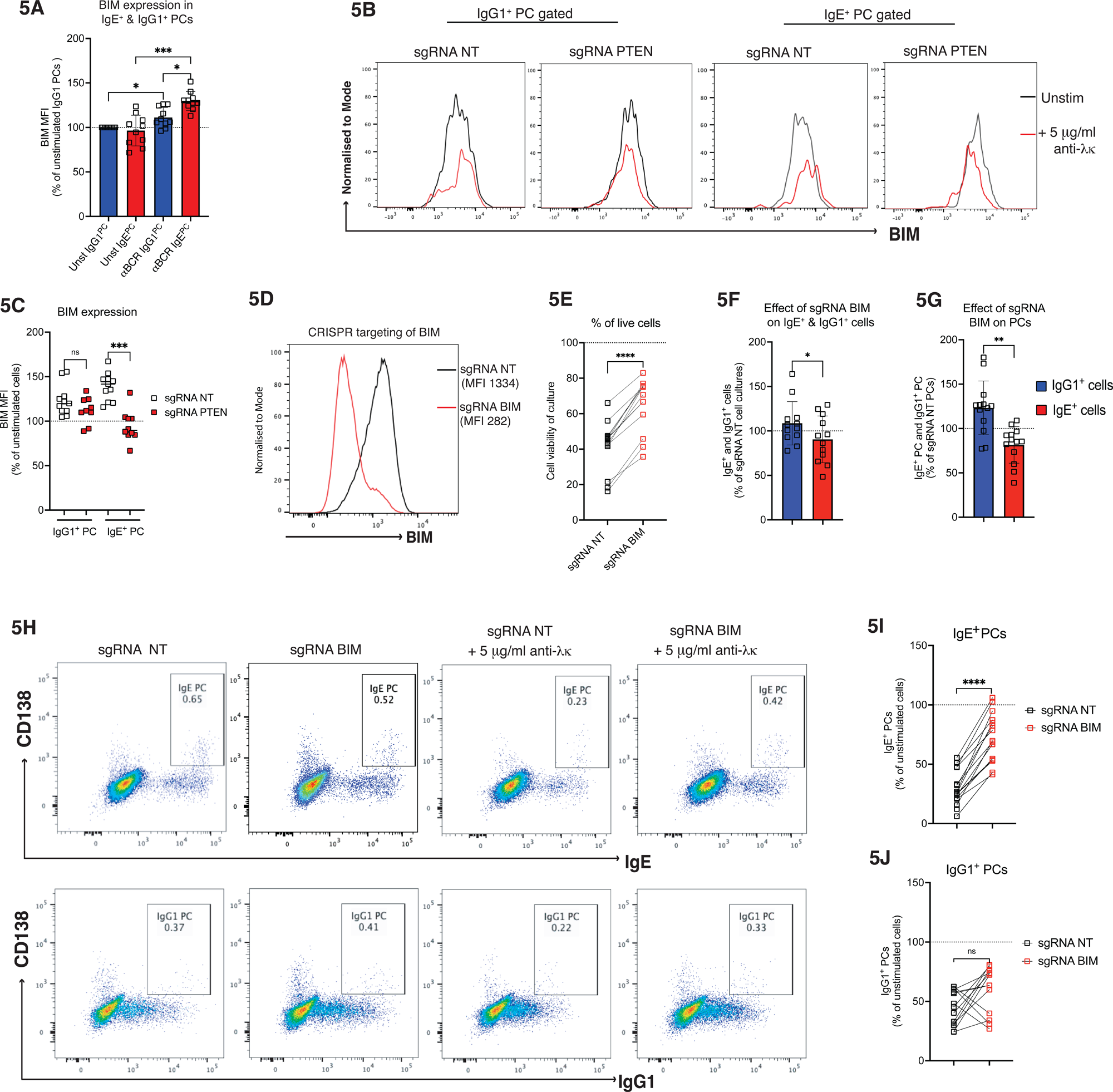
BIM upregulation mediates preferential BCR killing of IgE⁺ PCs. (A) BIM expression (MFI) in unstimulated and BCR-crosslinked (anti-λ/anti-κ F(ab′)₂) IgE⁺ and IgG1⁺ PCs. BIM induction is significantly greater in IgE⁺ PCs following BCR engagement. (B) Representative histograms of BIM expression in IgG1⁺ and IgE⁺ PCs from control sgRNA NT and sgRNA PTEN cultures, shown under unstimulated conditions (black line) or following 48h of BCR crosslinking (red line). Histograms are normalised to mode. (C) BIM expression in IgG1⁺ and IgE⁺ PCs in sgRNA NT and sgRNA PTEN cultures following BCR crosslinking. Data shown as a percentage of unstimulated culture. (D) Representative histogram confirming efficient CRISPR/Cas9-mediated targeting of BIM (sgRNA BIM) compared with control sgRNA NT. (E) Percentage of viable cells (eFleur 788 negative) in sgRNA NT and sgRNA BIM cultures. (F) Effect of BIM targeting on the proportions of IgE⁺ and IgG1⁺ cells in class-switching cultures, shown as a percentage of sgRNA NT cultures. (G) Effect of BIM targeting on IgE^+^ and IgG1^+^ PCs gated within the total IgE^+^ and IgG1^+^ cells. The data are shown as a percentage of sgRNA NT cell culture. (H) Representative flow cytometry plots showing IgE⁺ (top) and IgG1⁺ (bottom) PCs in sgRNA NT and sgRNA BIM cultures, in the presence or absence of BCR crosslinking. (I) IgE⁺ PCs and IgG1⁺ PCs (**J**) in sgRNA NT and sgRNA BIM cell cultures following BCR crosslinking. Data is shown as a percentage of unstimulated culture. Each symbol represents individual donors. The data indicate mean ± SD. Statistical significance was determined using one-way ANOVA with Tukey’s multiple comparison test (A) or paired two-tailed t-test with Welch’s correction (C, F, G, I, J). *P < 0.05, **P < 0.01, ***P < 0.001, ****P < 0.0001; ns, not significant.

To directly test the role of BIM in BCR-mediating killing of IgE^+^ PCs, we targeted BIM using CRISPR/Cas9 at the onset of the culture. Compared with non-targeting sgRNA cultures, sgRNA targeting BIM reduced BIM expression (Figure 5D) and increased the overall cell viability of the cultures (Figure 5E). It also modestly reduced the proportion of IgE^+^ cells compared to IgG1^+^ cells (Figure 5F). Interestingly, BIM targeting also led to a significant increase in the proportion of IgG1^+^ PCs but not IgE^+^ PCs (Figure 5G).

To determine whether BIM contributes to the BCR-mediated killing of IgE^+^ and IgG1^+^ PCs, class-switched cells were cultured for an additional 48 h in the presence or absence of BCR crosslinking. BIM targeting led to a striking rescue of IgE^+^ PCs from BCR-mediated killing (Figure 5H and 5I). In contrast, the effect on IgG1^+^ PCs was modest and variable across experiments (Figure 5H and 5). Together, these data identify BIM as a key effector of BCR-mediated apoptosis in IgE^+^ PCs, acting downstream of elevated PTEN expression and reduced PI3K signaling to lower the threshold for apoptosis specifically in IgE^+^ PCs.

### Increased JNK signaling contributes to enhanced BIM expression and apoptosis in IgE⁺ PCs following BCR crosslinking

Previous studies have reported that IgE^+^ PCs exhibit elevated expression of genes associated with endoplasmic reticulum (ER) stress and protein production, which is consistent with their fivefold higher antibody secretion rates compared with IgG1^+^ PCs (Vecchione et al., 2024; Miranda-Waldetario et al., 2025). Sustained ER stress can promote apoptosis, in part through the activation of the JNK signaling pathway (Dhanasekaran and Reddy, 2008; Lee et al., 2011; Kaneto et al., 2005; Win et al., 2014; Rosati et al., 2010). In several systems, the PI3K-Akt and JNK signaling can have opposing effects on apoptosis (Aikin et al., 2004; Kim et al., 2001; Wang et al., 2007; Song et al., 2016), with PTEN acting as a modulator of this balance (Zhang et al., 2007). We therefore investigated whether JNK signaling is involved in the preferential killing of IgE^+^ PCs by BCR crosslinking. IgE^+^ PBs and PCs had modestly higher basal levels of JNK phosphorylation compared to their IgG1^+^ counterparts (Figure 6A and S6A). Consistent with the above data showing impaired BCR signaling in IgE^+^ GC-like B cells, BCR crosslinking induced higher JNK phosphorylation in IgG1^+^ GC-like B cells than in IgE^+^ GC-like B cells (Figure 6B and S6A). However, JNK phosphorylation was similar between IgE^+^ PBs and IgG1^+^ PBs, and significantly higher in IgE^+^ PCs compared to IgG1^+^ PCs after BCR crosslinking (Figure 6B and S6A). Thus, BCR crosslinking induces higher JNK activation in IgE^+^ PCs than in IgG1^+^ PCs.

**Figure 6.**
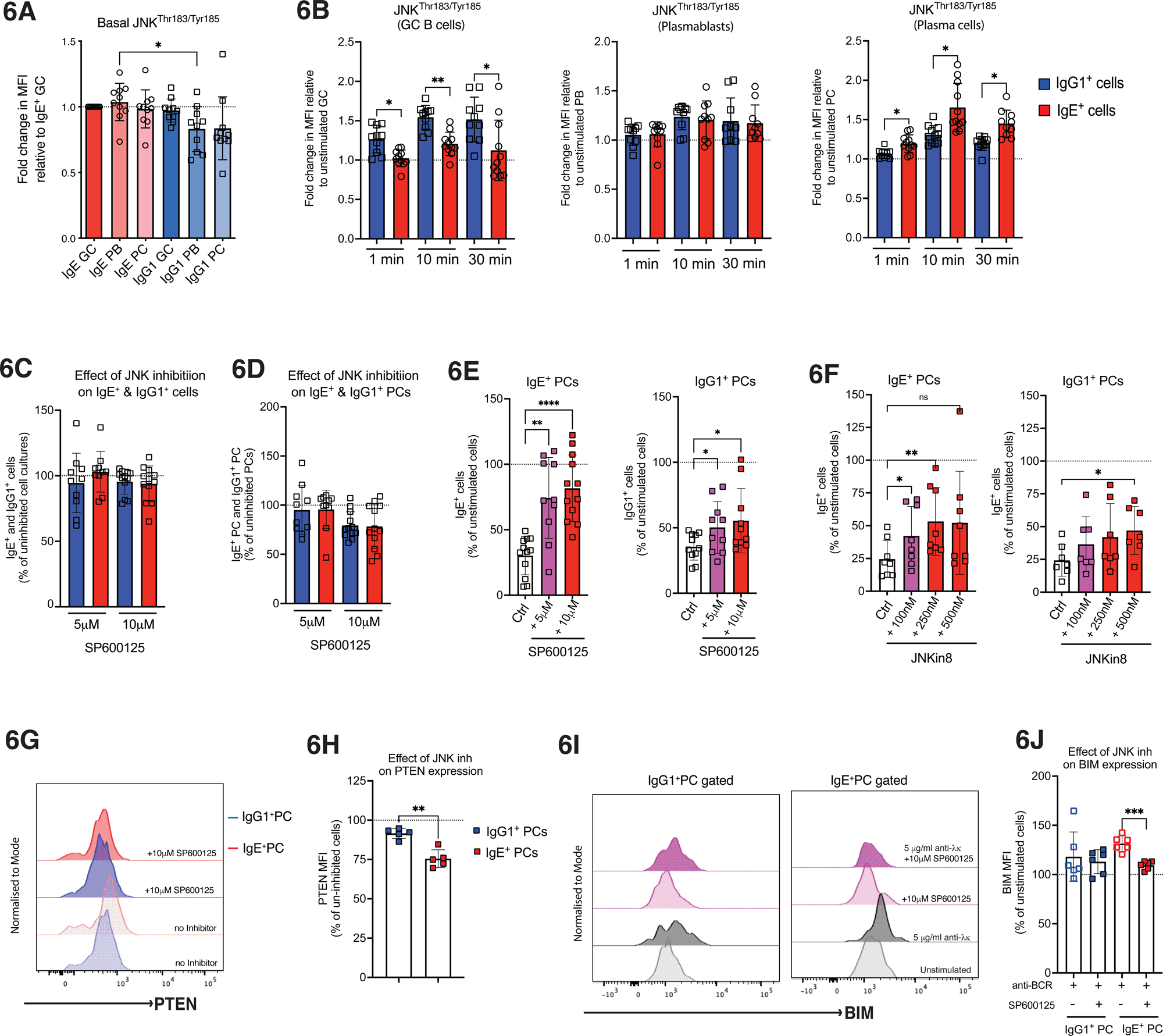
JNK signaling promotes PTEN expression, BIM induction, and BCR induced apoptosis in IgE⁺ PCs. (A) Basal phosphorylation of JNK (Thr183/Tyr185) in IgE⁺ and IgG1⁺ GC-like B cells, plasmablasts (PBs), and plasma cells (PCs), expressed as fold change in mean fluorescence intensity (MFI) relative to IgE⁺ GC-like B cells. (B) JNK phosphorylation following BCR crosslinking in GC-like B cells, PBs, and PCs at the indicated time points (1, 10, and 30 min), shown as fold change in MFI relative to unstimulated GC-like B cells, PBs, and PCs. (C) Effect of JNK inhibition (SP600125; 5 μM and 10 μM) on the proportions of IgE⁺ and IgG1⁺ cells in class-switch cultures. The data are shown as a percentage of uninhibited cell culture. (D) Effect of JNK inhibition on IgE⁺ and IgG1⁺ PCs gated within the total IgE^+^ and IgG1^+^ cells. The data are shown as a percentage of uninhibited cell culture. (E) Effect of SP600125 on BCR-mediated killing of IgE⁺ PCs (left) and IgG1⁺ PCs (right). Cells were cultured for 48 h with IL-4 and anti-CD40 in the presence or absence of 5 μg/mL F(ab’)_2_ anti-λ/κ and inhibitor. Ctrl = IL4+ anti-CD40 with DMSO. Data is shown as a percentage of the unstimulated cell culture. (F) Validation of JNK-dependent effects on BCR mediated killing using the irreversible JNK inhibitor JNK-IN-8 (100–500 nM) on IgE⁺ PCs (left) and IgG1⁺ PCs (right). (G) Representative histograms showing PTEN expression in IgE⁺ and IgG1⁺ PCs after 48 h of culture with IL4 and anti-CD40 and in the presence or absence of 10 μM SP600125. (H) PTEN expression in IgE⁺ and IgG1⁺ PCs following JNK inhibition, shown as a percentage of uninhibited controls. (I) Representative histograms of BIM expression in IgG1⁺ PCs (left) and IgE⁺ PCs (right) under unstimulated conditions or following 48 h of BCR crosslinking, with or without JNK inhibition. (J) BIM expression in IgE⁺ and IgG1⁺ PCs following BCR crosslinking in the presence or absence of SP600125, shown as a percentage of unstimulated culture controls. Data are pooled from independent experiments (symbols represent individual donors). Bars indicate mean ± SD. Statistical significance was determined using one-way ANOVA with Tukey’s multiple comparison test (B, E, F) or paired two-tailed t-test with Welch’s correction (H, J). *P < 0.05, **P < 0.01, ***P < 0.001, ****P < 0.0001; ns, not significant.

To assess the functional contribution of this enhanced JNK signaling to cell death, day 10 class-switched cells were cultured for a further 48 h with JNK inhibitor SP600125 (5 μM and 10 μM), in the presence or absence of BCR crosslinking. SP600125 did not significantly change the proportions of IgE^+^ and IgG1^+^ cells in the absence of BCR crosslinking (Figure 6C and 6D). However, following BCR crosslinking, both concentrations of SP600125 rescued the IgE^+^ PCs from the BCR-mediated killing (Figure 6E and S6B). A significant rescue in killing was also observed in IgG1^+^ PCs, but this was more modest (Figure 6E and S6C). Similar effects were observed using JNK-IN-8, an irreversible covalent inhibitor of JNK (Zhang et al., 2012) (Figure 6F), confirming that JNK activity is required for the BCR-mediated killing, particularly in IgE^+^ PCs.

Notably, inhibition of JNK also significantly reduced PTEN expression in IgE^+^ PCs, with a more limited effect in IgG1^+^ PCs (Figure 6G and 6H). This suggests that JNK signaling contributes to PTEN expression in IgE^+^ PCs, thus opposing the PI3K dependent survival in these cells. Consistently, we found that JNK inhibition prevented the upregulation of BIM in IgE^+^ PCs following BCR crosslinking, whereas BIM expression in IgG1^+^ PCs was largely unaffected (Figure 6I and 6J).

Together, these data support a model in which enhanced JNK signaling in IgE^+^ PCs sustains PTEN expression, which limits PI3K survival signals and promotes a Ca^2+^- and BIM-dependent apoptosis following BCR crosslinking.

## Discussion

BCR signaling plays a key role in regulating B cell development, survival and differentiation, yet its impact on PC biology is not very well understood. In this study, we demonstrate that BCR crosslinking preferentially induces apoptosis in human IgE⁺ PCs due to an imbalance between pro-survival and stress signaling pathways. Elevated PTEN expression in IgE⁺ PCs constrains their PI3K/Akt signaling, lowering the apoptotic threshold by enabling enhanced induction of the pro-apoptotic BIM following BCR crosslinking. In parallel, JNK signaling acts as a critical amplifier of this pro-apoptotic programme, revealing a coordinated JNK/PTEN/BIM axis that selectively predisposes IgE⁺ PCs to cell death.

We show that crosslinking of the IgE BCR, using either therapeutic antibodies, such as omalizumab and EMPD-targeting antibodies, or antibodies crosslinking the BCR via binding to the Ig light chains, reduces IgE production by selectively killing IgE⁺ cells, with the most pronounced effect on IgE⁺ PCs. Enhanced apoptosis, either spontaneous or induced by BCR binding to antigen, has been shown in mouse IgE^+^ PCs (Newman and Tolar, 2021; Wade-Vallance et al., 2023). The mouse IgE^+^ PCs express enhanced levels of the IgE BCR compared to IgE^+^ GC-like cells and compared to IgG^+^ PCs, which downmodulate their BCRs (Yang et al., 2012; Wade-Vallance et al., 2023). Indeed, the recently generated IgG1^+^ PCs still express detectable levels of the BCR but about 10-fold lower than those on IgE^+^ PCs (Wade-Vallance et al., 2023). In contrast, IgM^+^ and IgA^+^ PCs maintain BCR surface expression upon differentiation (Wade-Vallance et al., 2023; Pinto et al., 2013; Blanc et al., 2016). We show that in human PCs, BCR surface expression is similar between IgE^+^ and IgG^+^ PCs. Potentially, this may reflect the usage of the primate-specific long isoform of the mIgE, which reduces surface IgE BCR expression in IgE^+^ PCs (Vanshylla et al., 2018; Ramadani et al., 2017). This observation reinforces the conclusion that the pro-apoptotic activity is a specific feature of the IgE BCR in IgE⁺ PCs, which lose protection from apoptosis once IL-4 and CD40 signaling wane with the induction of the PC differentiation (Ramadani et al., 2019).

Importantly, these findings identify a previously underappreciated cellular mechanism through which therapeutic antibodies, such as omalizumab, may suppress IgE production by directly targeting IgE^+^ PCs and inducing their apoptosis. In the case of omalizumab, binding likely occurs preferentially to soluble IgE (Holgate, 2014; Guntern and Eggel, 2020), and its access to mIgE on PCs may be limited. However, prolonged omalizumab therapy can lead to sustained clinical benefits even after treatment discontinuation (Foti Randazzese et al., 2025; Humbert et al., 2022; Deschildre et al., 2019; Nopp et al., 2007), rasing the possibility that omalizumab leads to the depletion of IgE^+^ PCs *in vivo*. Indeed, long-term follow-up studies have reported that a substantial proportion of patients maintain improved asthma control several years after cessation of omalizumab therapy (Foti Randazzese et al., 2025; Humbert et al., 2022; Deschildre et al., 2019; Nopp et al., 2007). Potentially, an early treatment neutralises circulating IgE, followed by progressive depletion of IgE⁺ PCs as mIgE becomes accessible.

We show indeed that anti-IgE CεmX 4B12 antibody, which targets the EMPD of mIgE, is more efficient at killing IgE⁺ PCs than omalizumab. This likely reflects improved accessibility to the mIgE, as CεmX 4B12 does not bind to soluble IgE (Chen et al., 2010). Notably, we observed that IgE⁺ GC-like B cells are relatively resistant to both omalizumab and anti-IgE CεmX 4B12 targeting, suggesting that these recently switched IgE^+^ B cells, together with type 2 (CD23^+^IgG1^+^) memory B cells (Koenig et al., 2024; Ota et al., 2024), may act as a reservoirs that replenishes IgE⁺ PCs following killing. These observations have important implications for anti-IgE therapy. Clinical trials using quilizumab monotherapy, an antibody targeting the EMPD of mIgE, failed to show meaningful clinical benefits in patients with allergic asthma and chronic spontaneous urticaria (Harris et al., 2016a; b). Thus, therapeutic strategies combining EMPD targeting antibodies with omalizumab could accelerate and enhance the depletion of IgE^+^ PCs while simultaneously neutralising serum IgE, potentially reducing the need for prolonged treatment. In addition, considering the potential for the new generation of IgE^+^ PCs, combining anti-IgE therapy with dupilumab, which blocks IL-4 receptor alpha and inhibits de novo class switching to IgE (Hamilton et al., 2021; Guttman-Yassky et al., 2019; Dekkers et al., 2023; Wenzel et al., 2013), may provide a more comprehensive approach to targeting IgE production. Emerging clinical data suggest that such dual biologic strategies may provide improved disease control (Arasu et al., 2022; Silva et al., 2025).

A central finding of our study is that the mechanism underlying the preferential sensitivity of human IgE^+^ PC to BCR-mediated apoptosis is fundamentally different to that established in mice. Previous work in mice linked this apoptotic sensitivity of IgE^+^ PC to higher surface BCR expression and enhanced proximal IgE BCR signaling, specifically to increased Syk and PLCγ2 activation (Wade-Vallance et al., 2023). In contrast, our data demonstrate that the apoptotic sensitivity of human IgE^+^ PC cannot be explained by these mechanisms as we find that in human cells, surface BCR expression and phosphorylation of Syk and PLCγ2 is similar between IgE⁺ and IgG1⁺ PCs, and that BCR signaling through Syk and PLCγ2 is required for BCR-mediated killing of both IgE⁺ and IgG1⁺ PCs. This is further reinforced by the observation that mouse IgM⁺ PCs exhibit similar proximal BCR signaling characteristics to IgE⁺ PCs, including comparable activation of Syk and PLCγ2, yet do not display the same susceptibility to BCR-induced apoptosis (Wade-Vallance et al., 2023). These observations indicate that the preferential killing of IgE^+^ PCs by BCR crosslinking is not determined by the strength of proximal signaling but rather by differences in how these signals are integrated downstream of the BCR.

Our data identify attenuated PI3K/Akt signaling as the key factor of this differential sensitivity to BCR crosslinking. Indeed, IgE⁺ PCs showed reduced Akt phosphorylation following BCR crosslinking, which was consistent with the elevated expression of PTEN in these cells. In B cells, increased PTEN has been shown to function as a critical checkpoint that tunes the balance between survival and apoptosis during the antigen-driven selection of immature B cells, where increased PTEN activity favours BCR-induced apoptosis over survival signaling (Cheng et al., 2009; Wang et al., 2022). The elevated levels of PTEN in IgE^+^ PCs thus likely function as a signaling switch that constrains survival while promoting pro-apoptotic signals downstream of the IgE BCR as in the case of peripheral anergic/autoreactive B cells (Smith et al., 2019). In this context, our finding that IgE⁺ PCs selectively upregulate PTEN suggests that PTEN is an important regulator of IgE production. The central role of the PI3K signaling in these cells is also supported by the PI3K p110δ inhibition experiments, which preferentially enhance BIM expression and cell death in IgE^+^ PCs. In contrast, CRISPR/Cas9 targeting of PTEN preferentially rescues IgE⁺ PCs, but not IgG1^+^ and IgM^+^ PCs, from BCR-mediated apoptosis, further supporting the conclusion that elevated PTEN imposes a selective constraint on survival signaling in IgE⁺ PCs following BCR crosslinking.

Consistent with impaired PI3K signaling, we show that BCR crosslinking induces a stronger upregulation of the pro-apoptotic protein BIM in IgE⁺ PCs than in IgG1⁺ PCs. This induction is dependent on PTEN, as its targeting prevents BIM upregulation in IgE^+^ PCs. Moreover, direct targeting of BIM substantially rescues IgE⁺ PCs from BCR-mediated apoptosis, identifying BIM as a key downstream effector linking attenuated survival signaling to apoptosis in IgE⁺ PCs. We further show that Ca²⁺ signaling contributes to the induction of apoptosis, as inhibition of Ca²⁺ flux reduces BCR-mediated killing, with a greater protective effect in IgE⁺ PCs than in IgG1⁺ PCs. Together, these findings indicate that Ca²⁺ dependent and BIM-mediated apoptotic pathways are more active in IgE⁺ PCs following BCR crosslinking. Mechanistically, we find that IgE^+^ PCs have elevated JNK activation and inhibition of JNK significantly rescues IgE⁺ PCs from BCR-mediated apoptosis. Furthermore, we show that JNK inhibition reduces PTEN expression and prevents BIM induction, indicating that JNK reinforces the pro-apoptotic state of IgE^+^ PCs following BCR crosslinking. Given that JNK is a well-established effector of stress and Ca²⁺dependent signaling (Dhanasekaran and Reddy, 2008; Lee et al., 2011; Kaneto et al., 2005; Win et al., 2014; Rosati et al., 2010), these findings support a model in which JNK sustains elevated PTEN expression, thereby constraining PI3K/Akt survival signaling and promoting BIM dependent apoptosis.

The origin of enhanced JNK activity and increased PTEN levels in IgE^+^ PCs remains to be determined. One possibility is that they originate from chronic, autonomous signaling from the IgE BCR in the absence of co-stimulatory signals such as CD40 and IL-4R. Another possibility is that they relate to enhanced ER stress. A distinctive feature of IgE⁺ PCs is the elevated expression of genes associated with protein synthesis and ER stress, and and higher antibody secretory activity (Vecchione et al., 2024; Miranda-Waldetario et al., 2025). The ER stress may be directly promoted by strong interactions of IgE BCR with ER chaperones and stress sensors (Obayashi et al., 2025). The ER plays a central role in Ca²⁺ homeostasis and protein folding (Chen et al., 2023; Makio et al., 2024; Krebs et al., 2015), and perturbations in these processes trigger the unfolded protein response (UPR). While UPR is physiologically part of the PC differentiation and initially promotes cell survival, when it is severe or sustained, UPR transitions to a pro-apoptotic programme (Gass et al., 2002; Wu and Kaufman, 2006; Grootjans et al., 2016; Ricci et al., 2021; Puthalakath et al., 2007). Notably, this programme involves the JNK pathway. In B cells, ER stress has also been linked to BCR-mediated apoptosis following crosslinking (Yan et al., 2008) suggesting that BCR signaling can converge with stress responses to promote apoptosis. In this context, the increased ER stress and JNK activation in IgE⁺ PCs may lower the threshold for apoptosis upon BCR crosslinking, in part by amplifying Ca²⁺ flux and downstream mitochondrial apoptosis pathways.

Together, our data support a model in which the outcome of BCR signaling is determined by the balance between survival and stress pathways. In IgE⁺ PCs, elevated PTEN expression attenuates PI3K signaling, allowing JNK and Ca²⁺dependent signaling pathways to dominate, leading to enhanced BIM induction and apoptosis following BCR crosslinking. These findings not only refine our understanding of IgE biology but also provide a mechanistic basis for therapeutic strategies aimed at eliminating IgE^+^ PCs, with implications for improving the efficacy of anti-IgE therapies.

## Materials and methods

### Ethics

Following full informed written consent, tonsils were obtained from patients undergoing routine tonsillectomies for recurrent tonsillitis or airway obstruction at the Guy’s and St Thomas’s NHS Fundation Trust (REC number 08/H0804/94) and at UCL Hospitals NHS Trust (REC number 23/LO/0869).

### Isolation of B cells and class switching cultures

Human B cells were isolated from fresh tonsils using 2-aminoethylisothiouronium bromide-treated sheep red blood cells (TCS Biosciences Ltd) as previously described (Ramadani et al., 2015, 2017). The B cells were then further enriched by magnetic cell sorting using the MojoSort Human pan B Cell Isolation Kit according to the manufacturer’s instructions (BioLegend). The purified tonsil B cells were cultured at 0.5 x10^6^ cells/mL in complete culture media, containing RPMI 1640 (without L-glutamine, Lonza) supplemented with penicillin (100 IU/mL, Invitrogen), streptomycin (100 mg/mL, Invitrogen) and glutamine (2 mM, Invitrogen) and 10% Foetal Calf Serum (FBS, Hyclone; Perbio Biosciences). To induce class switching to IgE, the cultures were supplemented with IL-4 (400 IU/ml; R&D Europe Systems Ltd and anti-CD40 antibody (1 μg/mL; G28.5;BioXCell), Transferrin (35 µg/mL) (Sigma-Aldrich) and insulin (5 µg/mL) (Sigma-Aldrich). The cells were incubated for 12 days at 37°C with 5% CO2. The class switched cells were then harvested, washed by centrifugation at 105 g for 10 min, and resuspended at 0.5 x10^6^ cells/mL in complete culture media, with or without IL4 and anti-CD40. Cells were then stimulated for up to 48 h with anti-IgE antibodies (5 μg/mL; anti-CεmX a20, anti-CεmX 4B12(Chen et al., 2010) or omalizumab) and either F(ab’)_2_ anti-κ (5 μg/mL) or F(ab’)_2_ anti-λ (5 μg/mL) alone or both F(ab’)_2_ anti-κ (5 μg/mL) and anti-λ (5 μg/mL).

### Culture of YK6 expressing CD40 ligand and BAFF

An immortalized tonsil FDC-like feeder cell, termed YK6, expressing CD40Lg and BAFF were a kind gift of Daniel J. Hodson (University of Cambridge)(Caeser et al., 2019). YK6-CD40Lg-BAFF cells were expanded in T175 cell culture flasks with complete advanced culture media, which contained Advanced RPMI media (Gibco) supplemented with 10% heat-inactivated FBS (Sigma), 100X GlutaMAX (Gibco), 100X HEPES Buffer (Gibco), 100X Minimum Essential Medium Non-Essential Amino Acid solution (Gibco), 100X Penicillin-Streptomycin (Gibco), 50 μM 2-mercaptoethanol (Gibco). The adherent cells were typically trypsinised when at 80-100% confluency. Cells were than irradiated with a dose of 40 Gy and either seeded in 6 well plates at 10 x 10^4^ YK6-CD40Lg-BAFF cells per well or stored at -80 C until further use.

### Single guide RNA (sgRNA) design

CRISPR sgRNAs were designed using Synthego and Integrated DNA Technologies (IDT) sgRNA design tools. sgRNAs with the highest predicted on-target and lowest off-target activity were ordered from Synthego and resuspended in TE buffer (Synthego) at 30μM. A non-targeting gecko sgRNA sequence was used as the negative control. Primers were designed and PCR-performed on extracted DNA to create amplicons that included the site of the predicted dsDNA break. Amplicons were sequenced using Sanger sequencing and the sgRNA that caused the highest percentage of indels was selected. The targeting success was then confirmed by flow cytometry. The selected sgRNAs are detailed in supplementary table 1.

### CRISPR of human tonsil B cells

Alt-R^TM^ S.p. Cas9 Nuclease V3 (IDT, 1081059) and sgRNAs were mixed at a 1:4 molar ratio and incubated at 23C for 30 min to form ribonucleoprotein (RNP) complexes. During this incubation, 1 x 10^6^ of purified tonsil B cells per reaction were washed with D-PBS and then resuspended in 15 ul per reaction with P3 Nucleofector solution from the P3 Primary Cell 4D-Nucleofector Kit (Lonza). Resuspended cells were then mixed with the RNP solution and transferred to the Nucleofector strip and electroporated using the 4D Lonza Nucleofector system (program EO-115). Following electroporation, 80 ul of warmed culture medium was added to the Nucleofector strip wells and the cells were transferred to a 6 well plate pre seeded with 10 x 10^4^ of irradiated YK6-CD40Lg-BAFF cells per well in complete advanced culture media supplemented with 400 IU/ml of IL4 (Preprotech). On day 7 of the culture, the cells were harvested and plated onto freshly seeded YK6-CD40Lg-BAFF cells and kept in culture until day 12. In the case of PTEN, we also first initiated class switching to IgE by culturing tonsil B cells for 7 days with 10 x 10^4^ YK6 YK6-CD40Lg-BAFF cells in complete advanced culture media supplemented with 400 IU/ml of IL4 (Preprotech). On day 7 of the cultures, the cells were harvested, washed and CRISPR gene targeted as above. The electroporated cells were than cultured for another 5 days and then used for follow up analysis or stimulated with both F(ab’)_2_ anti-κ (5 μg/mL) and anti-λ (5 μg/mL) for 48 h.

### Flow cytometry

Cultured cells were harvested and washed with D-PBS by centrifugation at 105 g for 10 min. The pelleted cells were resuspended in D-PBS and stained with a fixable viability dye eFluor780 (1/1000) for 20 min on ice. Cells were then washed and incubated for 10 mins with FACS buffer containing FcR block (Miltenyi Biotec) followed by staining with antibodies against surface markers on ice for 15 min. Cells were then washed with FACS buffer and fixed with 2% prewarmed paraformaldehyde (PFA) for 10 mins at 37°C, and permeabilized with PBS containing 0.5% saponin and 0.5% Triton x-100 for 20 mins at room temperature in the dark. The permeabilised cells were washed with FACS buffer and stained with antibodies against intracellular markers for 30 mins in the dark (see table for a list of antibodies used). Cells were then washed and resuspended in 200 µl FACS buffer and acquired using BD LSR Fortessa. The data were analysed using FlowJo software (Tree Star Inc, USA).

### Phosphoflow

Day 12 class switching cultures containing IgE^+^ and IgG1^+^ cells were harvested and pelleted by centrifugation at 105 g for 10 min. The cells were washed once with D-PBS and then resuspended at 2 x 10^6^ cells per ml of RPMI without L-glutamine and 2% FBS. Cells were rested for 2 h at 37C in a 5% CO2 incubator. Following the rest period, cells were stained with a fixable viability dye eFluor780 (1/1000) for 10 min at 37C. After the incubation, the cells were pelleted and resuspended at 1x10^6^ cells per 200 μl of RPMI without L-glutamine and stimulated for 1, 10 or 30 min with F(ab’)_2_ anti-κ (10 ug/ml) and F(ab’)_2_ anti-λ (10 ug/ml) antibodies. The BCR stimulation was stopped by the addition of prewarmed paraformaldehyde (final concentration 2%) and incubation for 10 min at 37C. The fixed cells were washed twice with D-PBS (800 g for 5 min) and permeabilized by slow addition of 1 ml of ice-cold methanol followed by incubation on ice for 20 min. The permeabilised cells were either stained immediately or stored for up to two weeks at -20°C until further analysis. Prior to staining, methanol was removed by washing the cells twice with ice cold D-PBS (800 g for 5 min) and rehydrating them with FACS buffer for 1 h at room temperature. Cells were then washed and incubated for 10 mins with FACS buffer containing FcR block (Miltenyi Biotec) followed by staining with antibodies against CD138, IgE, IgG1, and phosphorylated signaling proteins PLCg2, Syk, Akt and JNK (summarised in supplemental table 2). After staining, cells were washed, resuspended in FACS buffer and acquired using BD LSR Fortessa. Data were analysed using FlowJo software (Tree Star Inc).

### Caspase-3 activation

Day 12 class switching cultures were harvested, washed and resuspended in complete advanced culture media at 0.5 x 10^6^ cells per 1 ml. The cells were than stimulated with 10 ug/ml of F(ab’)_2_ anti-κ and anti-λ antibodies for 1 h at 37C in a 5% CO2 incubator. 10 min before the end of 1 h incubation the cells were stained with a fixable viability dye eFluor780 at a 1/1000 and gently mixed with a pipette. The cells were then washed with D-PBS and fixed for 10 min with 2% prewarmed paraformaldehyde at 37C. Following this, cells were permeabilised with 1 ml of ice-cold methanol and incubated for 20 min in ice. The permealised cells were either stained immediately as above or stored for up to two weeks at -20°C until further analysis.

### Total IgE secretion

The amounts of total IgE secreted in supernatants of cultures after 48 h of stimulations with anti-CεmX (a20 and 4B12) and omalizumab were measured using Phadia ImmunoCAP-100 (Thermo Fisher Scientific/ Phadia), according to the manufacturer’s instructions.

## Acknowledgements

We thank the IIT flow cytometry facilities for technical assistance. We are grateful to the patients and the ENT surgical teams of Guy’s & St Thomas’s and UCL NHS Fundation Trusts. We also thank Dr David Fear (King’s College London) and Dr Lizzy Rosser (University College London) for their support with the collection of tonsils used in this research. We are also grateful to Prof Hannah Gould (King’s College London) for early support of the project and Dr Jiun-Bo Chen and Prof. Tse Wen Chang (Academia Sinica, Taiwan) for the anti-IgE EMPD antibodies. This study was supported by an Asthma UK career development award to F.R. (AUK-CDA-2019-412), Guy’s and St Thomas’ Charity (R170502), and a Wellcome Trust Investigator Award to P.T. (223196/Z/21/Z). S. W. T. C. was supported by an MRC DTP training programme (MR/W006774/1). Graphical abstract figure was created with BioRender.com. For the purpose of Open Access, the authors have applied a CC BY public copyright licence to any Author Accepted Manuscript version arising from this submission.

## Author contributions

F.R. and P.T. designed the experiments. F.R., H.T., S.W.T. C., C. T. and L. O-L. performed experiments. F.R. and P.T. supervised the research. F.R. and P.T. analysed the data and wrote the paper.

