## Supplemental Material for "Altered PI3K-PTEN balance promotes preferential killing of human IgE^+^ plasma cells by BCR crosslinking"

### Supplemental Figures

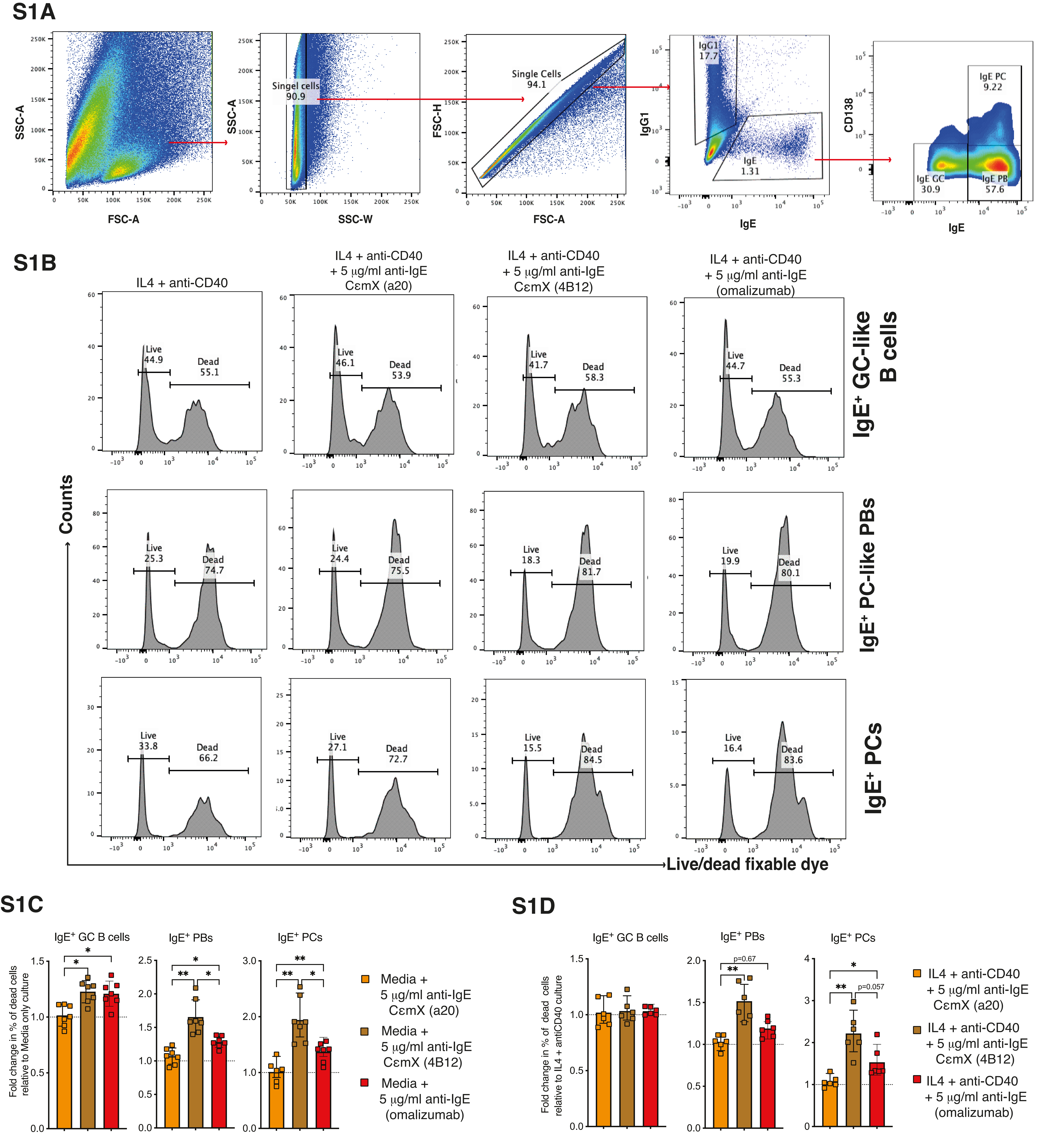
S1A
SSC-A
FSC-A
Singel cells 90.9
SSC-W
FSC-H
Single Cells 94.1
IgG1 17.7
IgE 1.31
IgG1
IgE
CD138
IgE PC 9.22
IgE GC 30.9
IgE PB 57.6
S1B
IL4 + anti-CD40
IL4 + anti-CD40 + 5 µg/ml anti-IgE CεmX (a20)
IL4 + anti-CD40 + 5 µg/ml anti-IgE CεmX (4B12)
IL4 + anti-CD40 + 5 µg/ml anti-IgE (omalizumab)
IgE⁺ GC-like B cells
IgE⁺ PC-like PBs
IgE⁺ PCs
Counts
Live/dead fixable dye
Live 44.9
Dead 55.1
Live 46.1
Dead 53.9
Live 41.7
Dead 58.3
Live 44.7
Dead 55.3
Live 25.3
Dead 74.7
Live 24.4
Dead 75.5
Live 18.3
Dead 81.7
Live 19.9
Dead 80.1
Live 33.8
Dead 66.2
Live 27.1
Dead 72.7
Live 15.5
Dead 84.5
Live 16.4
Dead 83.6
S1C
IgE⁺ GC B cells
IgE⁺ PBs
IgE⁺ PCs
Fold change in % of dead cells relative to Media only culture
Media + 5 µg/ml anti-IgE CεmX (a20)
Media + 5 µg/ml anti-IgE CεmX (4B12)
Media + 5 µg/ml anti-IgE (omalizumab)
S1D
IgE⁺ GC B cells
IgE⁺ PBs
IgE⁺ PCs
Fold change in % of dead cells relative to IL4 + antiCD40 culture
p=0.67
p=0.057
IL4 + anti-CD40 + 5 µg/ml anti-IgE CεmX (a20)
IL4 + anti-CD40 + 5 µg/ml anti-IgE CεmX (4B12)
IL4 + anti-CD40 + 5 µg/ml anti-IgE (omalizumab)

**Figure S1. Gating strategy and viability of IgE$^+$ cell subsets following BCR crosslinking.**

(A) Class switched cell cultures were first gated based on forward scatter (FSC-A) and side scatter (SSC-A), followed by exclusion of doublets using SSC-W and FSC-A parameters. Single cells were then gated into IgE$^+$ cells that were further gated based on CD138 expression into GC-like B cells (IgE$^{low}$CD138$^-$), plasmablasts (PBs; IgE$^{hi}$CD138$^-$), and plasma cells (PCs; IgE$^{hi}$CD138$^+$). Numbers indicate the percentage of cells within each gate.

(B) Representative histograms showing live/dead staining with a fixable viability dye in IgE$^+$ GC-like B cells (top row), PBs (middle row), and PCs (bottom row) cultured with IL-4 and anti-CD40, in the absence or presence of 5 µg/mL anti-IgE antibodies (CεmX a20, CεmX 4B12, or omalizumab). Percentages of live and dead cells are indicated.

(C) Cell viability in IgE$^+$ GC-like B cells, PBs, and PCs cultured in media alone and stimulated with anti-IgE antibodies. Data are presented as fold change relative to unstimulated controls.

(D) Cell viability in IgE$^+$ GC-like B cells, PBs, and PCs cultured in IL-4 and anti-CD40 and stimulated with anti-IgE antibodies. Data are presented as fold change relative to unstimulated IL-4 and anti-CD40 controls.

Data represent independent donors, with each symbol indicating an individual culture. Bars show mean ± SD. Each symbol represents individual donors. The data indicate mean ± SD. Statistical significance was determined using one-way ANOVA with Tukey’s multiple comparison test (C, D). *$P < 0.05$, **$P < 0.01$, ***$P < 0.001$, ns, not significant.

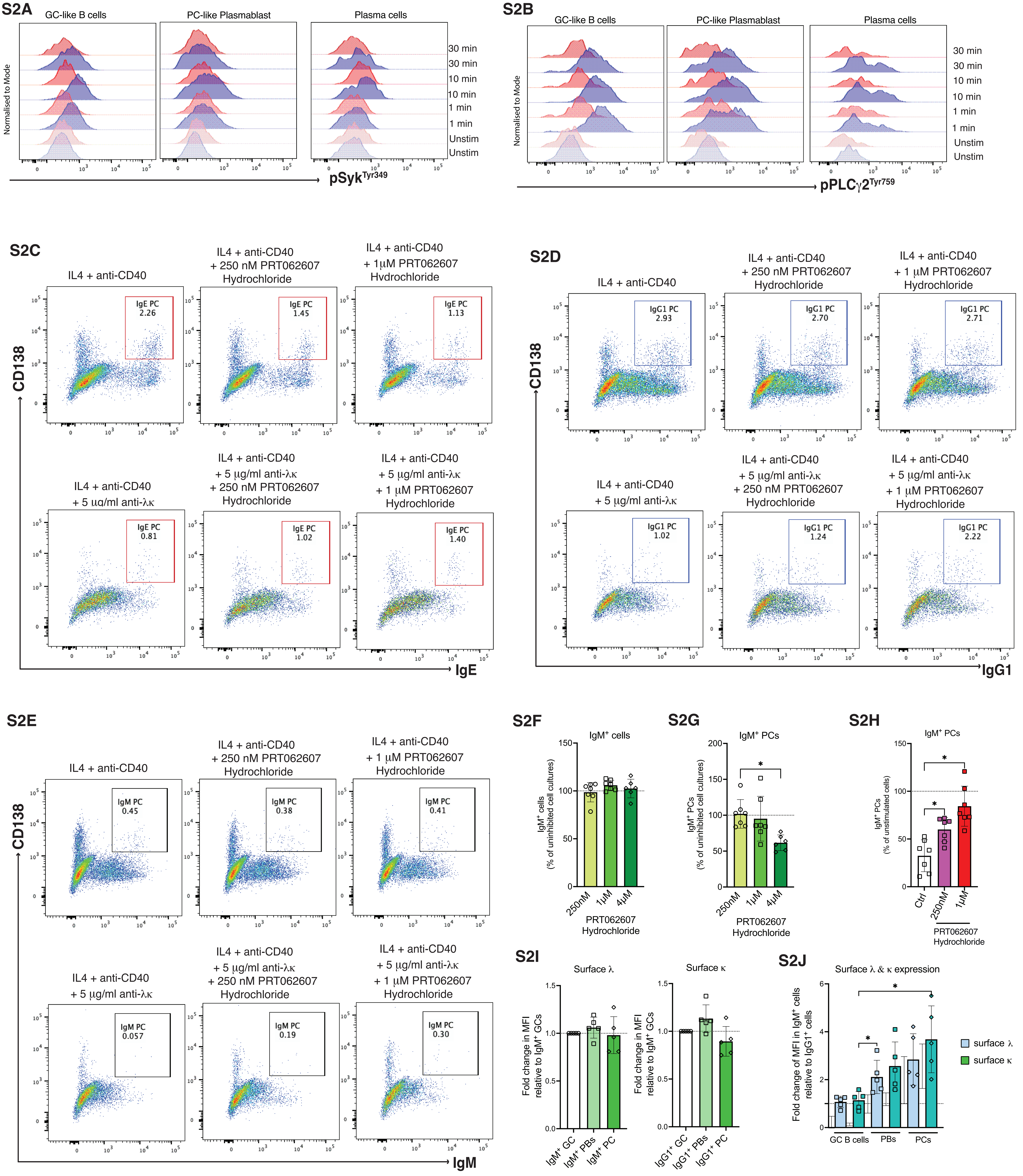

S2A
GC-like B cells
PC-like Plasmablast
Plasma cells
Normalised to Mode
30 min
30 min
10 min
10 min
1 min
1 min
Unstim
Unstim
pSyk Tyr349
S2B
GC-like B cells
PC-like Plasmablast
Plasma cells
Normalised to Mode
pPLCγ2 Tyr759
S2C
IL4 + anti-CD40
IL4 + anti-CD40 + 250 nM PRT062607 Hydrochloride
IL4 + anti-CD40 + 1μM PRT062607 Hydrochloride
IgE PC 2.26
IgE PC 1.45
IgE PC 1.13
IL4 + anti-CD40 + 5 μg/ml anti-λκ
IL4 + anti-CD40 + 5 μg/ml anti-λκ + 250 nM PRT062607 Hydrochloride
IL4 + anti-CD40 + 5 μg/ml anti-λκ + 1 μM PRT062607 Hydrochloride
IgE PC 0.81
IgE PC 1.02
IgE PC 1.40
CD138
IgE
S2D
IL4 + anti-CD40
IL4 + anti-CD40 + 250 nM PRT062607 Hydrochloride
IL4 + anti-CD40 + 1 μM PRT062607 Hydrochloride
IgG1 PC 2.93
IgG1 PC 2.70
IgG1 PC 2.71
IL4 + anti-CD40 + 5 μg/ml anti-λκ
IL4 + anti-CD40 + 5 μg/ml anti-λκ + 250 nM PRT062607 Hydrochloride
IL4 + anti-CD40 + 5 μg/ml anti-λκ + 1 μM PRT062607 Hydrochloride
IgG1 PC 1.02
IgG1 PC 1.24
IgG1 PC 2.22
CD138
IgG1
S2E
IL4 + anti-CD40
IL4 + anti-CD40 + 250 nM PRT062607 Hydrochloride
IL4 + anti-CD40 + 1 μM PRT062607 Hydrochloride
IgM PC 0.45
IgM PC 0.38
IgM PC 0.41
IL4 + anti-CD40 + 5 μg/ml anti-λκ
IL4 + anti-CD40 + 5 μg/ml anti-λκ + 250 nM PRT062607 Hydrochloride
IL4 + anti-CD40 + 5 μg/ml anti-λκ + 1 μM PRT062607 Hydrochloride
IgM PC 0.057
IgM PC 0.19
IgM PC 0.30
CD138
IgM
S2F
IgM+ cells
IgM+ cells (% of uninhibited cell cultures)
250nM
1μM
4μM
PRT062607 Hydrochloride
S2G
IgM+ PCs
IgM+ PCs (% of uninhibited cell cultures)
250nM
1μM
4μM
PRT062607 Hydrochloride
S2H
IgM+ PCs
IgM+ PCs (% of unstimulated cells)
Ctrl
250nM
1μM
PRT062607 Hydrochloride
S2I
Surface λ
Surface κ
Fold change in MFI relative to IgM+ GCs
Fold change in MFI relative to IgM+ GCs
IgM+ GC
IgM+ PBs
IgM+ PC
IgG1+ GC
IgG1+ PBs
IgG1+ PC
S2J
Surface λ & κ expression
Fold change of MFI in IgM+ cells relative to IgG1+ cells
GC B cells
PBs
PCs
surface λ
surface κ

**Figure S2. BCR signaling and effects of Syk inhibition across PC subsets.**

(A–B) Representative histograms showing phosphorylation of proximal BCR signaling molecules following BCR crosslinking. Day 12 class-switched cultures were stimulated with anti-κ and anti-λ F(ab')$_2$ antibodies for the indicated times (1, 10 and 30 min). Phosphorylation of Syk (Syk$^{Tyr348}$**)** (A) and PLCγ2 (PLCγ2$^{Tyr759}$) (B) was assessed in IgE$^+$ and IgG1$^+$ GC-like B cells, PBs, and PCs. Histograms are normalised to mode, with unstimulated controls shown for comparison.

(C–D) Representative flow cytometry plots showing the effect of Syk inhibition (PRT062607 hydrochloride; 250 nM or 1 μM) on the frequency of IgE$^+$ PCs (C) and IgG1$^+$ PCs **(**D) in IL-4 and anti-CD40 cultures, in the presence or absence of BCR crosslinking (anti-κ/λ). Numbers indicate the percentage of cells within the gated PC population.

(E) Representative flow cytometry plots showing the effect of Syk inhibition on IgM$^+$ PCs under the same culture and stimulation conditions as in (C–D).

(F) IgM$^+$ cell frequencies following inhibition with increasing concentrations of PRT062607. Data are expressed as a percentage of uninhibited cell cultures.

(G) Effect of Syk inhibition on IgM$^+$ PCs gated within the total IgM$^+$ cells. The data are shown as a percentage of uninhibited cell culture.

(H) IgM$^+$ PCs following 48 h of BCR crosslinking, with or without Syk inhibition. Ctrl = IL4 + anti-CD40 with DMSO. Data are expressed as a percentage of unstimulated controls.

(I) Quantification of surface λ and κ mean fluorescence intensity (MFI) across differentiation stages of IgM$^+$ cells made relative to the MFI of IgM$^+$ GC-like B cells.

(J) Comparison of total surface BCR (λ + κ) expression between IgM$^+$ and IgG1$^+$ cells. The data show the fold change in MFI of λ and κ within each IgM$^+$ cell population made relative to their respective IgG1$^+$ cell counterparts.

Data represent independent donors, with each symbol corresponding to an individual culture. Bars indicate mean ± SD. Statistical significance was determined using one-way ANOVA with Tukey's multiple comparison test (H).*$P < 0.05$.

S3A

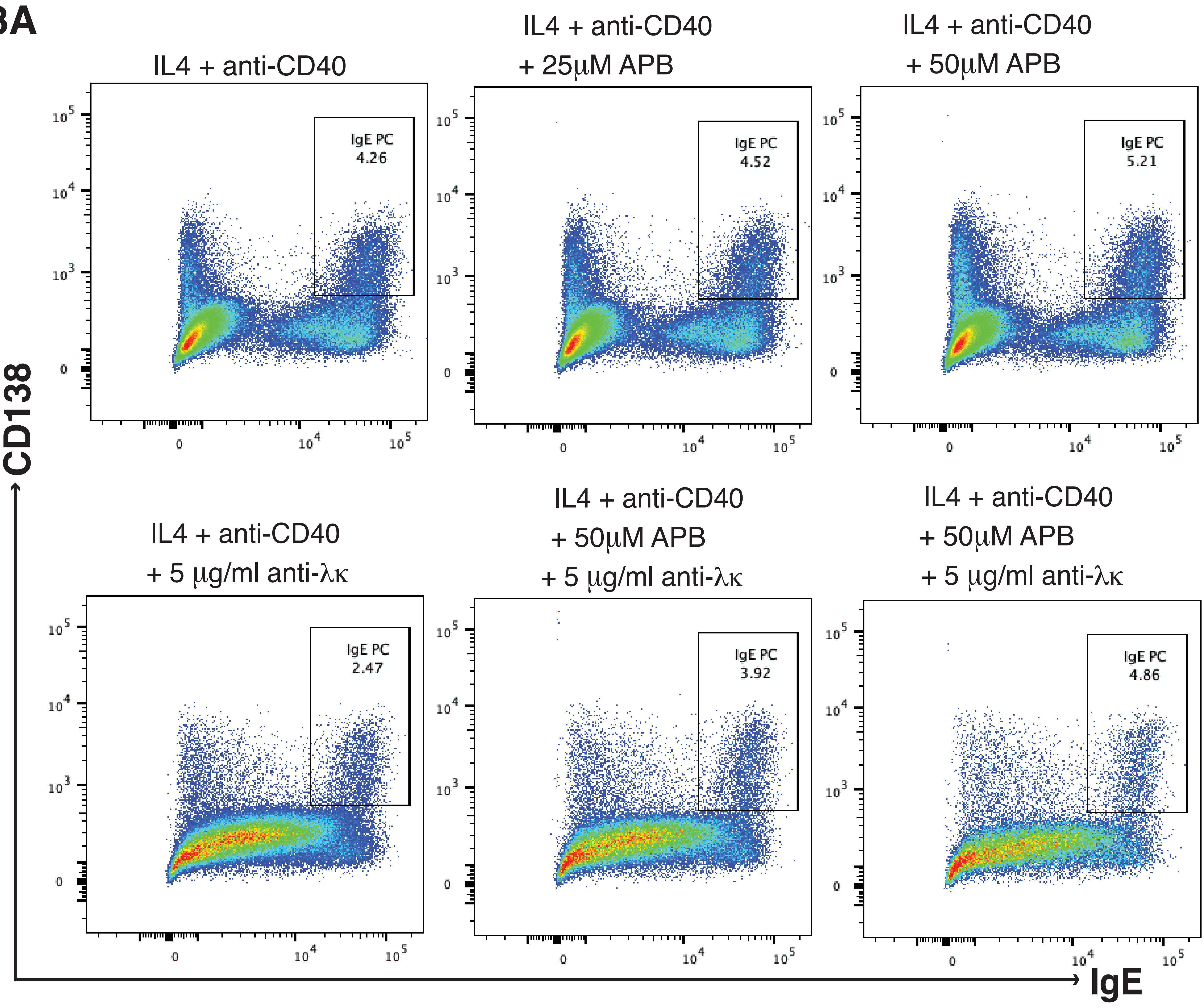


S3B

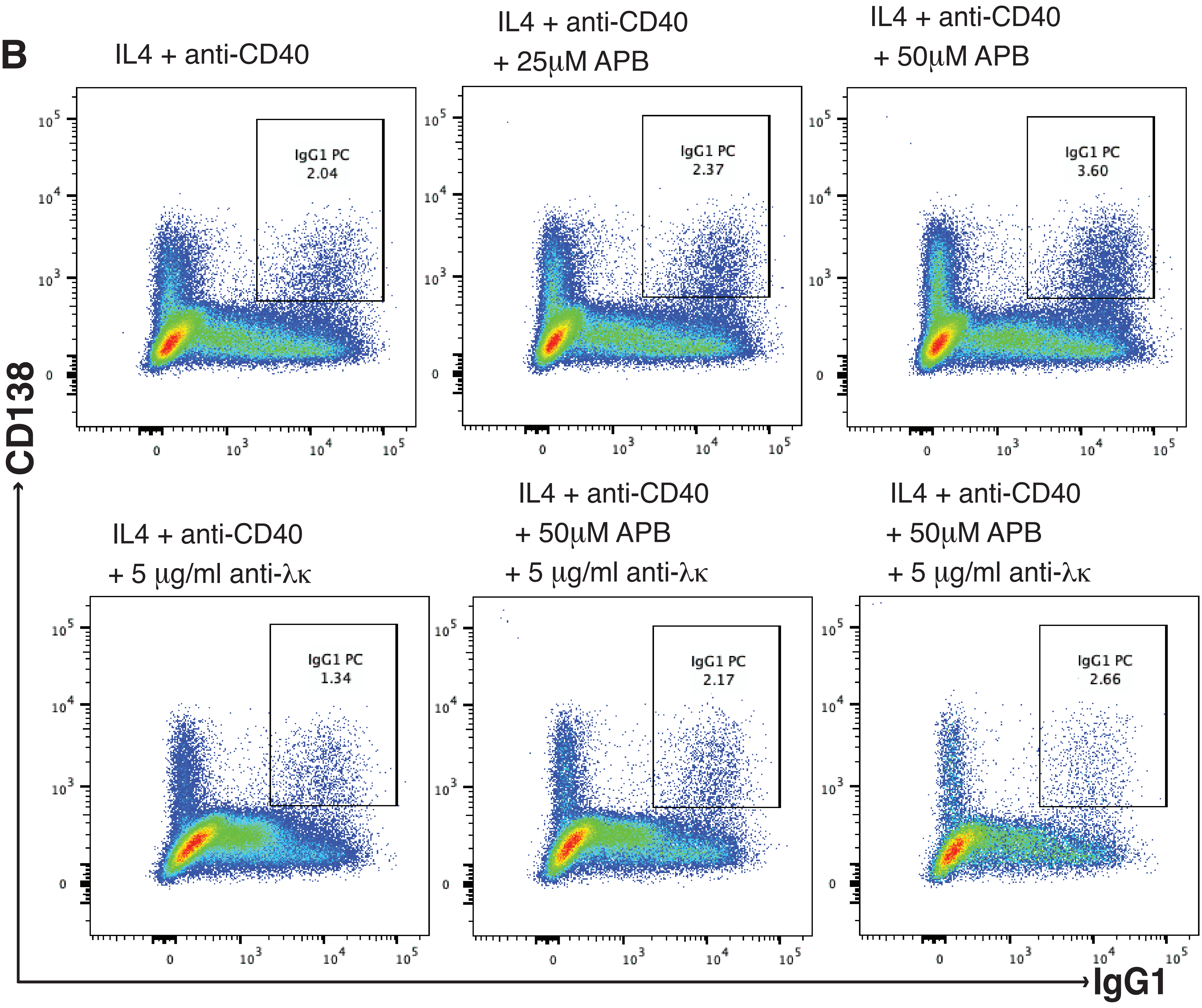


S3C

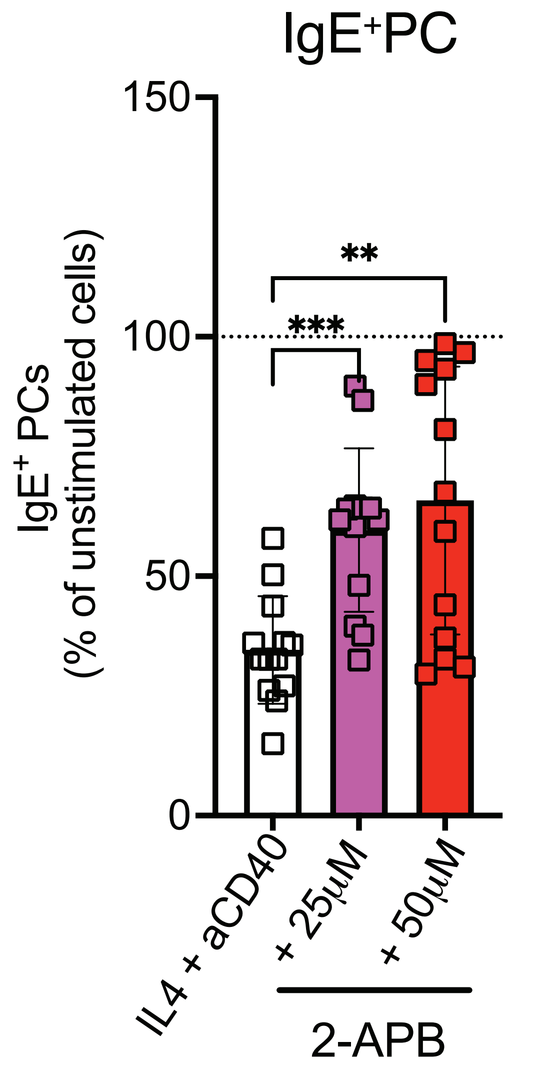


S3D

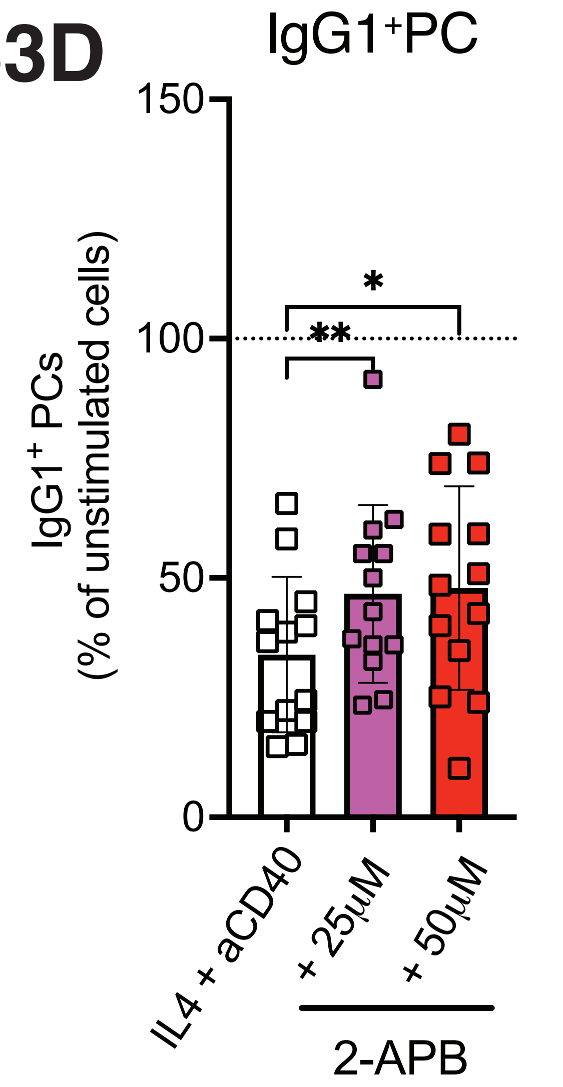

**Figure S3. Modulation of intracellular $Ca^{2+}$ signaling alters PC survival following BCR crosslinking.**

Representative flow cytometry plots showing the effect of 2-aminoethoxydiphenyl borate (2-APB; 25 μM or 50 μM) on the frequency of $IgE^+$ PCs (A) and $IgG1^+$ PCs (B). Class-switched cells were cultured for another 48 h in IL-4 and anti-CD40, in the absence or presence of BCR crosslinking (anti-κ/λ). Numbers indicate the percentage of cells within the gated PC population.

(C) $IgE^+$ PCs and $IgG1^+$ PCs (D) following 48 h of stimulation with 5 μg/ml of $F(ab')_2$ anti-λ/anti-κ in the presence of two different concentrations of 2-APB inhibitor. Data is shown as a percentage of the unstimulated cell culture.

Data represent independent donors, with each symbol corresponding to an individual culture. Bars indicate mean ± SD. Statistical significance was determined using one-way ANOVA with Tukey's multiple comparison test (C, D).$^*P < 0.05$, $^{**}P < 0.01$, $^{***}P < 0.001$.

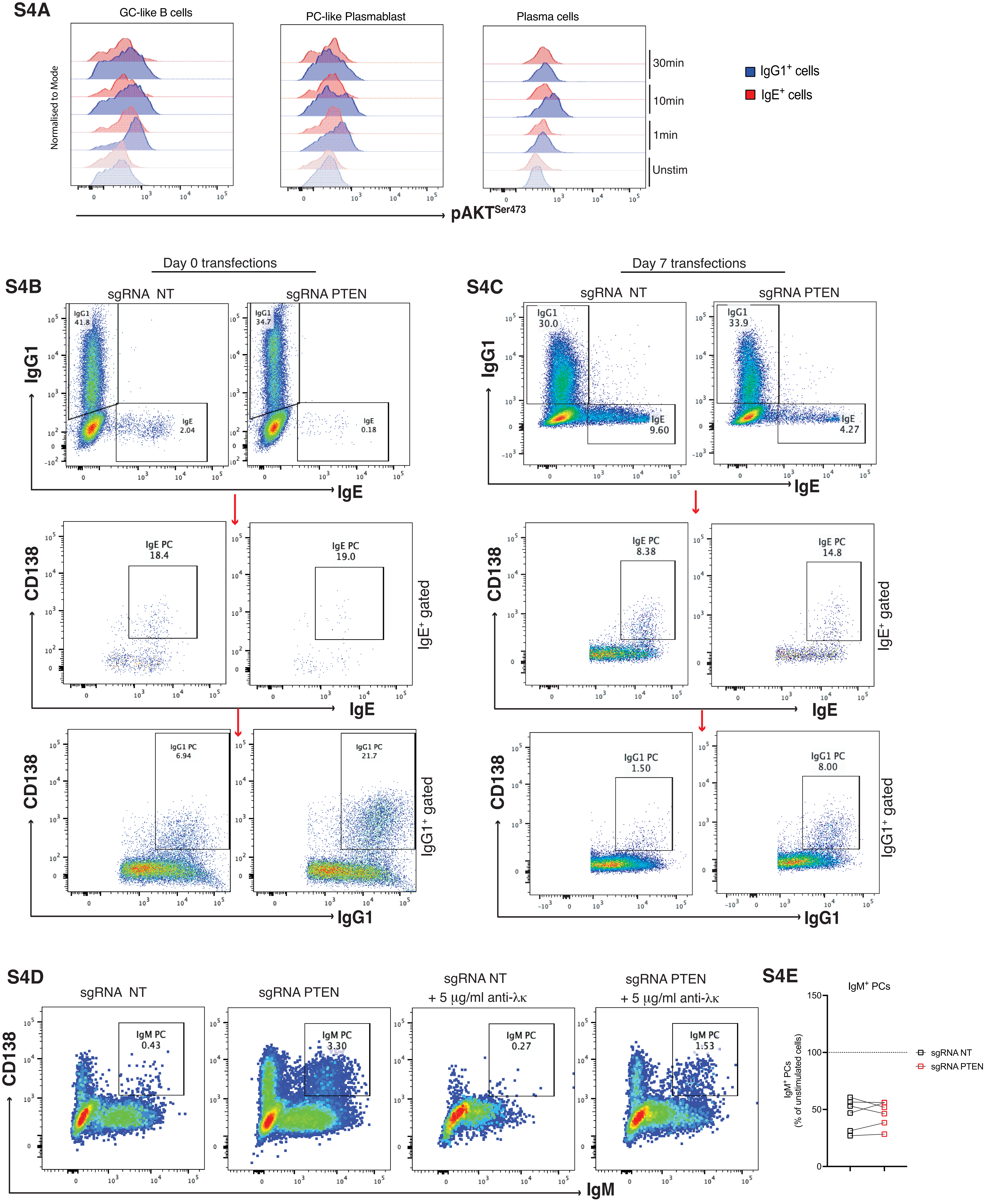

S4A
GC-like B cells
PC-like Plasmablast
Plasma cells
Normalised to Mode
30min
10min
1min
Unstim
IgG1+ cells
IgE+ cells
pAKTSer473
Day 0 transfections
Day 7 transfections
S4B
S4C
sgRNA NT
sgRNA PTEN
IgG1
IgE
IgG1 41.8
IgE 2.04
IgG1 34.7
IgE 0.18
IgG1 30.0
IgE 9.60
IgG1 33.9
IgE 4.27
CD138
IgE PC 18.4
IgE PC 19.0
IgE PC 8.38
IgE PC 14.8
IgE+ gated
IgG1 PC 6.94
IgG1 PC 21.7
IgG1 PC 1.50
IgG1 PC 8.00
IgG1+ gated
S4D
sgRNA NT + 5 µg/ml anti-λκ
sgRNA PTEN + 5 µg/ml anti-λκ
IgM PC 0.43
IgM PC 3.30
IgM PC 0.27
IgM PC 1.53
IgM
S4E
IgM+ PCs
IgM+ PCs (% of unstimulated cells)
150
100
50
0

**Figure S4. PTEN limits PI3K–Akt signaling and constrains PC survival and differentiation.**

(A) Representative histograms showing phosphorylation of Akt ($AKT^{Ser473}$) in $IgE^+$ (red) and $IgG1^+$ (blue) cells at different stages of differentiation (GC-like B cells, PC-like PBs and PCs). Cells were stimulated with anti-λ and anti-κ $F(ab')_2$ antibodies for the indicated times (1, 10 and 30 min) or left unstimulated. Data are normalised to mode.

(B) Representative flow cytometry plots of cultures electroporated on day 0 with sgRNA NT or sgRNA PTEN. Top panels show IgE versus IgG1 expression; middle panels show CD138 versus IgE ($IgE^+$ gate) and bottom panels show CD138 versus IgG1 ($IgG1^+$ gate).

(C) Representative flow cytometry plots of cultures electroporated on day 7 (after initiation of class switching) with sgRNA NT or sgRNA PTEN. Top panels show IgE versus IgG1 expression; middle panels show CD138 versus IgE within the gated $IgE^+$ cells, and bottom panels show CD138 versus IgG1 within the gated $IgG1^+$ cells.

(D) Representative flow cytometry plots of $IgM^+$ cells following PTEN targeting (day 7), cultured in the absence or presence of BCR crosslinking (5 µg/mL anti-λ/κ $F(ab')_2$). CD138 versus IgM plots identify $IgM^+$ PCs. PTEN

(E) $IgM^+$ PCs following BCR crosslinking, shown as a percentage of unstimulated controls, in sgRNA NT and sgRNA PTEN conditions. Each symbol represents an independent donor; lines connect paired samples.

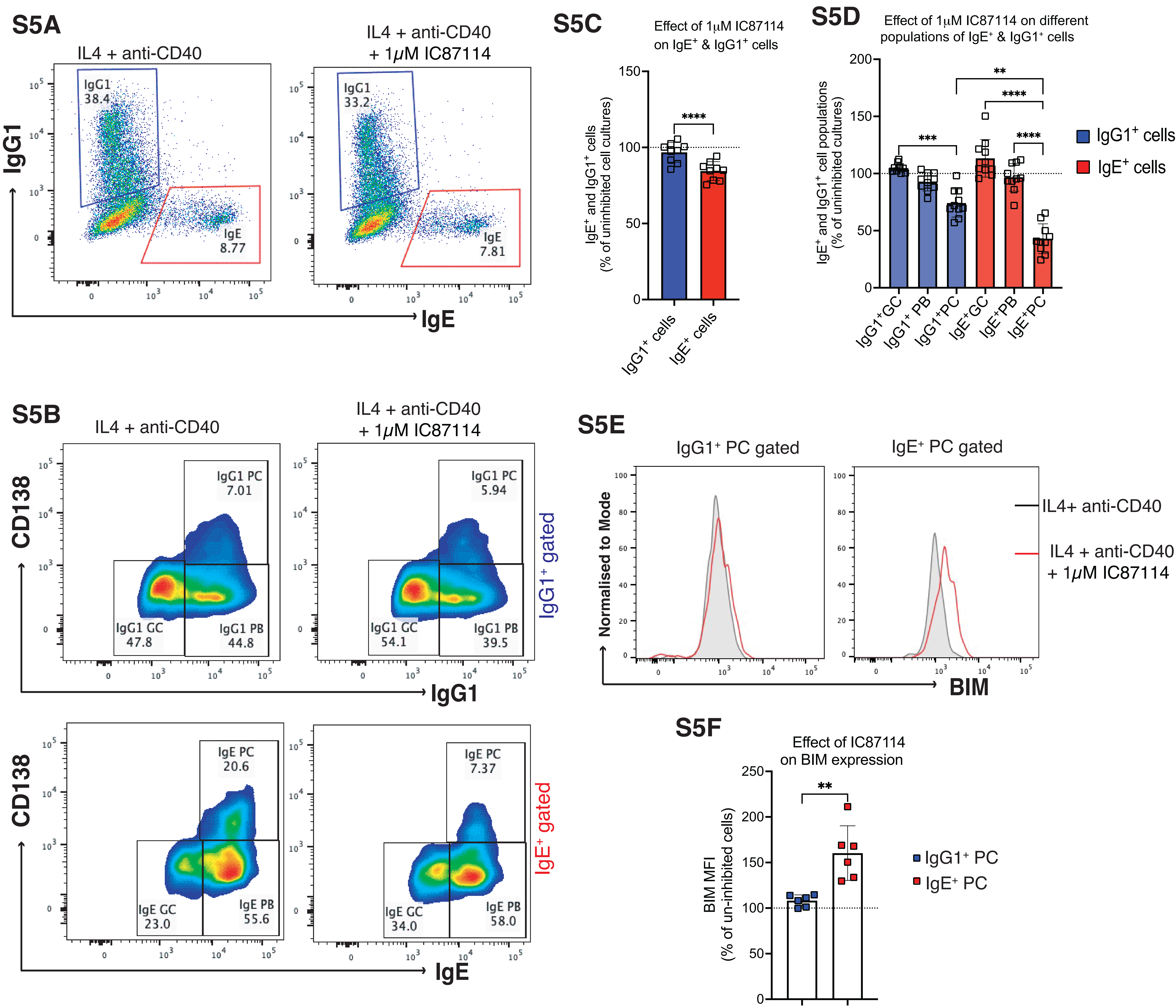
S5A
IL4 + anti-CD40
IL4 + anti-CD40
+ 1μM IC87114
IgG1
38.4
IgE
8.77
IgG1
33.2
IgE
7.81
IgG1
IgE
S5C
Effect of 1μM IC87114
on IgE⁺ & IgG1⁺ cells
IgE⁺ and IgG1⁺ cells
(% of uninhibited cell cultures)
****
IgG1⁺ cells
IgE⁺ cells
S5D
Effect of 1μM IC87114 on different
populations of IgE⁺ & IgG1⁺ cells
IgE⁺ and IgG1⁺ cell populations
(% of uninhibited cultures)
**
****
***
****
IgG1⁺GC
IgG1⁺ PB
IgG1⁺PC
IgE⁺GC
IgE⁺PB
IgE⁺PC
IgG1⁺ cells
IgE⁺ cells
S5B
IL4 + anti-CD40
IL4 + anti-CD40
+ 1μM IC87114
CD138
IgG1 PC
7.01
IgG1 GC
47.8
IgG1 PB
44.8
IgG1 PC
5.94
IgG1 GC
54.1
IgG1 PB
39.5
IgG1⁺ gated
IgG1
CD138
IgE PC
20.6
IgE GC
23.0
IgE PB
55.6
IgE PC
7.37
IgE GC
34.0
IgE PB
58.0
IgE⁺ gated
IgE
S5E
IgG1⁺ PC gated
IgE⁺ PC gated
Normalised to Mode
IL4+ anti-CD40
IL4 + anti-CD40
+ 1μM IC87114
BIM
S5F
Effect of IC87114
on BIM expression
BIM MFI
(% of un-inhibited cells)
**
IgG1⁺ PC
IgE⁺ PC

**Figure S5. PI3K p110δ inhibition selectively impairs IgE⁺ PC differentiation and enhances BIM expression.**

(A) Representative flow cytometry plots of IgE$^+$ and IgG1$^+$ cells cass-switched cells cultured for 48 h in IL-4 and anti-CD40, in the absence or presence of 1 μM IC87114. Numbers indicate the percentage of gated IgE$^+$ and IgG1$^+$ cells

IgE and IgG1 expression in day 12 class-switch cultures maintained with IL-4 and anti-CD40, in the absence or presence of the PI3K p110δ inhibitor IC87114 (1 μM). Gates indicate IgG1$^+$ (blue) and IgE$^+$ (red) populations.

(B) Representative flow cytometry plots showing the effect of 1 μM IC87114 on IgG1$^+$ (top panels) and IgE$^+$ PCs (bottom panels) gated within the total IgE$^+$ and IgG1$^+$ cells. Cultures were treated as in (A).

(C) Effect of PI3K p110δ (IC87114) on IgE$^+$ and IgG1$^+$ cells in the 48 h cultures. The data are shown as a percentage of uninhibited cell culture.

(D) Effect of PI3K p110δ on IgG1$^+$ and IgE$^+$ GC-like B cells, PBs and PCs. gated within IgG1$^+$ and IgE$^+$ cells. The data are shown as a percentage of uninhibited cell culture.

(E) Representative histograms of BIM expression in IgG1$^+$ and IgE$^+$ PCs cultured with IL-4 and anti-CD40, in the absence (grey filled) or presence (red line) of 1 μM IC87114. Data are normalised to mode.

(F) BIM MFI in IgG1$^+$ and IgE$^+$ PCs treated with 1 μM IC87114, shown as a percentage of uninhibited cell culture controls.

Data represent independent donors, with each symbol corresponding to an individual culture. Bars indicate mean ± SD. Statistical significance was determined using one-way ANOVA with Tukey's multiple comparison test (D) or paired two-tailed t-test with Welch's correction (C, F). *$P < 0.05$, **$P < 0.01$, ***$P < 0.001$.

S6A

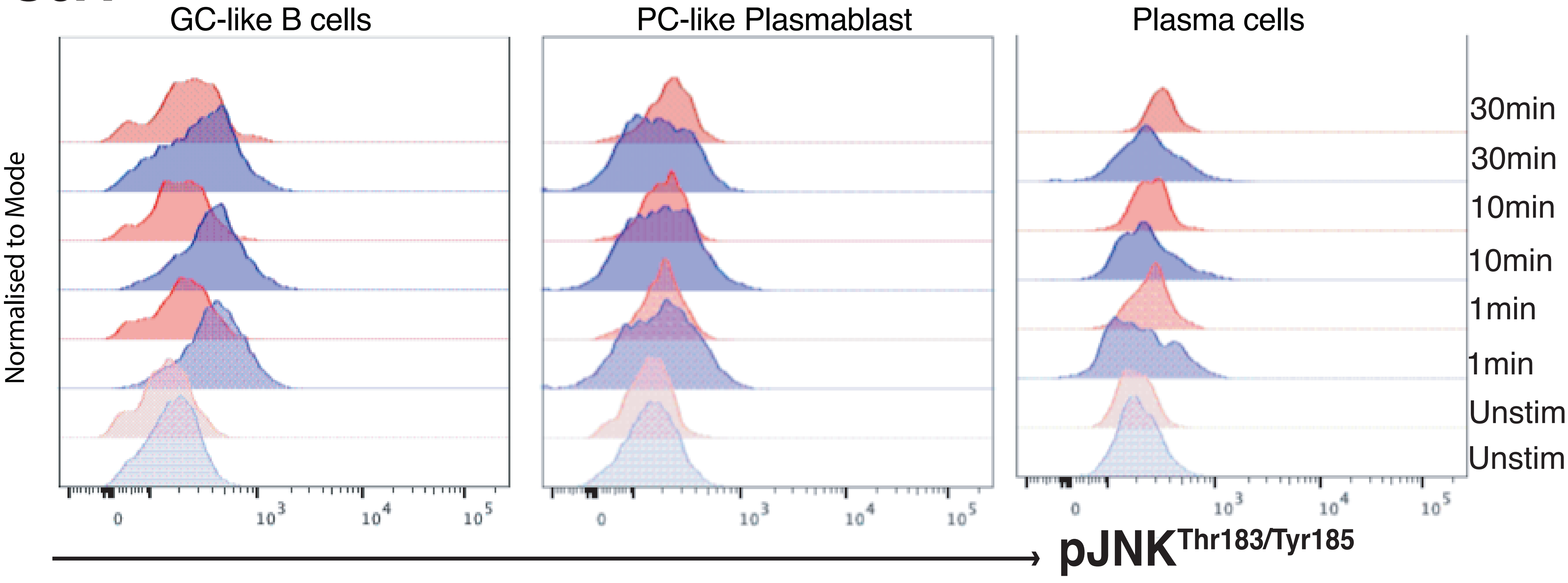

GC-like B cells
PC-like Plasmablast
Plasma cells
Normalised to Mode
30min
30min
10min
10min
1min
1min
Unstim
Unstim
pJNK Thr183/Tyr185


S6B

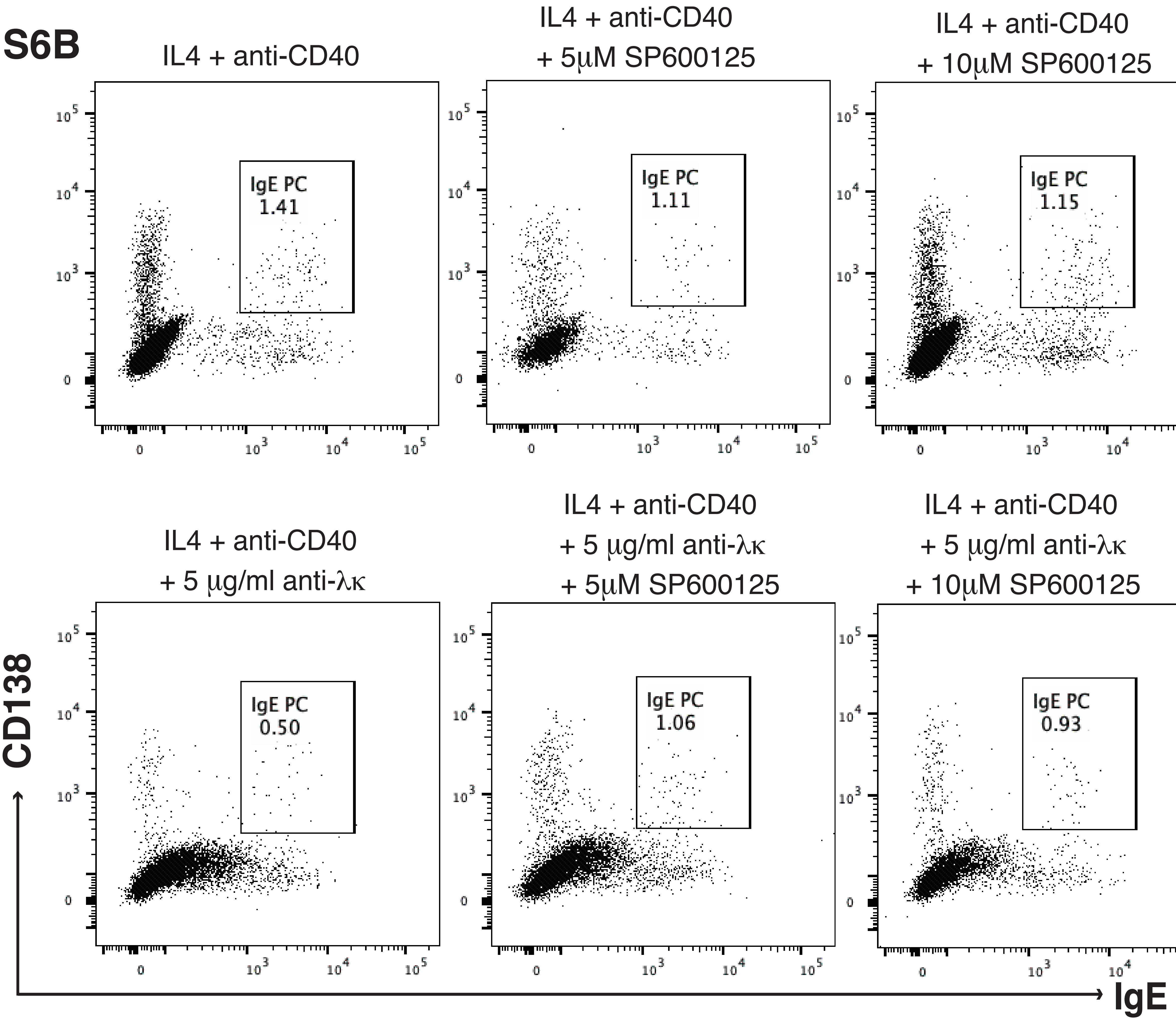

IL4 + anti-CD40
IgE PC 1.41
IL4 + anti-CD40 + 5µM SP600125
IgE PC 1.11
IL4 + anti-CD40 + 10µM SP600125
IgE PC 1.15
IL4 + anti-CD40 + 5 µg/ml anti-λκ
IgE PC 0.50
IL4 + anti-CD40 + 5 µg/ml anti-λκ + 5µM SP600125
IgE PC 1.06
IL4 + anti-CD40 + 5 µg/ml anti-λκ + 10µM SP600125
IgE PC 0.93
CD138
IgE


S6C

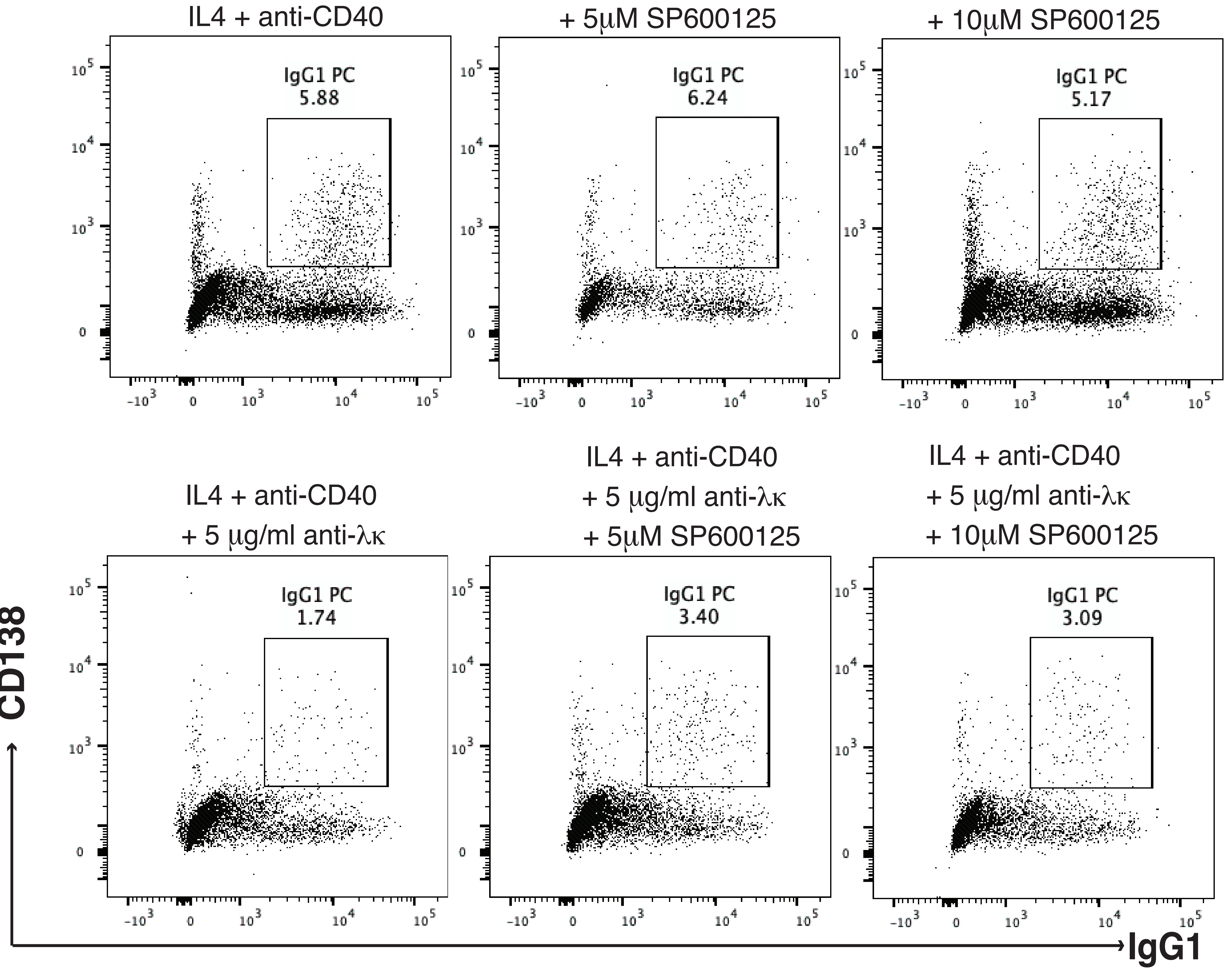

IL4 + anti-CD40
IgG1 PC 5.88
IL4 + anti-CD40 + 5µM SP600125
IgG1 PC 6.24
IL4 + anti-CD40 + 10µM SP600125
IgG1 PC 5.17
IL4 + anti-CD40 + 5 µg/ml anti-λκ
IgG1 PC 1.74
IL4 + anti-CD40 + 5 µg/ml anti-λκ + 5µM SP600125
IgG1 PC 3.40
IL4 + anti-CD40 + 5 µg/ml anti-λκ + 10µM SP600125
IgG1 PC 3.09
CD138
IgG1

**Figure S6. JNK signaling contributes to BCR killing of PCs, with a stronger effect in $IgE^+$ cells.**

(A) Representative histograms of JNK phosphorylation ($JNK^{Thr183/Tyr185}$) in $IgE^+$ (red) and $IgG1^+$ (blue) cells at different stages of differentiation (GC-like B cells, PC-like PBs, and PCs). Cells were stimulated with 10 μg/ml anti-λ and anti-κ $F(ab')_2$ antibodies for the indicated times (1, 10, and 30 min) or left unstimulated.

(B) Representative flow cytometry plots of $IgE^+$ PCs in IL-4 and anti-CD40 cultures treated with the JNK inhibitor SP600125 (5 μM or 10 μM), in the absence (top panels) or presence (bottom panels) of 5 μg/mL anti-λ/κ $F(ab')_2$.

(C) Representative flow cytometry plots of $IgG1^+$ PCs under the same conditions as in (B).

### Supplemental Tables

**Table S1**: sgRNA sequences to successfully target genes.

| Gene Target | sgRNA sequence |
|---|---|
| *Negative Control* | TGTAAACTGTCCAAGTAG |
| *Bcl2l11* | GTTCTGATGCAGCTTCCATG |
| *Pten* | GACTGGGAATAGTTACTCCC |
| *Plcγ2* | GAACUGAGUGCCAUAUAGGA |

**Table S2.** Antibodies used for flow cytometry staining.

| Antibody target | Clone | Company | Conjugate | Dilution |
|---|---|---|---|---|
| Active Caspase-3 | C92-605.rMAb | BD Biosciences | FITC | 1/50 |
| BIM | Y36 | Abcam | PE | 1/100-200 |
| | | | APC | 1/100-200 |
| CD19 | SJ25C1 | BD Biosciences | PE Cy7 | 1/25 |
| | HIB19 | | BUV 395 | 1/50 |
| CD138 | MI15 | Biolegend | BV 421 | 1/25 |
| Fixable Viability Dye | N/A | Life Technologies | eFluor 780 | 1/1500 |
| IgE | G7-2 | BD Biosciences | BUV 395 | 1/300 |
| | MHE-18 | Biolegend | APC | 1/300 |
| | | | PE Cy7 | 1/300 |
| IgG1 | IS11-12E4.23.20 | Miltenyi Biotec | PE | 1/600 |
| | | | APC | 1/1200 |
| IgM | MHM-88 | Biolegend | APC | 1/300 |
| | | | PE Cy7 | 1/300 |
| p-Syk (pY348) | I120-722 | BD Biosciences | Alexa Fluor 488 | 1/5 |
| | Moch1ct | eBioscience | PE | 1/20 |
| p-PLCγ2 (pY759) | REA341 | Miltenyi Biotec | PE-Vio 770 | 1/10 |
| PLCγ2 | K88-1161 | BD Biosciences | AF 647 | 1/20 |
| PTEN | REA270 | Miltenyi Biotec | PE | 1/10 |
| p-Akt (pS473) | REA359 | Miltenyi Biotec | APC | 1/10 |
| | A21001C | Biolegend | AF 647 | 1/20 |
| p-JNK (pT183/pY185) | N9-66 | BD Biosciences | PE | 1/5 |
| | | | APC | 1/5 |
| Kappa light chain | Polyclonal | Bio-Rad | PE | 1/50 |
| Lambda light chain | Polyclonal | Bio-Rad | FITC | 1/100 |
